# Morphological features of the domestic house cricket (*Acheta domesticus*) for translational aging studies

**DOI:** 10.1101/2025.03.26.645610

**Authors:** Gerald Yu Liao, Sherwin Dai, Elizabeth Bae, Swastik Singh, Jenna Klug, Christina Pettan-Brewer, Warren Ladiges

**Affiliations:** Department of Comparative Medicine, University of Washington School of Medicine, Seattle, WA, USA

**Keywords:** aging, morphology, house crickets, standardized husbandry, locomotion, sensory adaptation, geroscience

## Abstract

Aging alters morphology and locomotor function in diverse organisms, yet standardized model systems for studying these changes remain limited to a relatively few species. Here, we present a comprehensive analysis of age- and sex-dependent morphological variations in house crickets (*Acheta domesticus*), integrating refined husbandry protocols to enhance reproducibility and translational relevance. We observed progressive increases in body weight, length, and appendage dimensions with age, with pronounced sexual dimorphism emerging post-maturity. Structural adaptations, including increased femoral volume and cross-sectional area, suggest compensatory mechanisms for age-related declines in muscle efficiency, while reduced hind leg-to-body length ratios indicate potential biomechanical constraints on locomotion. Antennal growth patterns highlight prolonged sensory investment, potentially offsetting declining mobility in aging individuals. To ensure data consistency, we implemented a standardized husbandry framework incorporating self-determined photoperiods, co-housing both sexes, and controlled diet and hydration strategies. Our results underscore the necessity of harmonizing environmental conditions in gerontological research, as variations in lighting, substrate availability, and microbiome exposure may significantly impact physiological resilience and behavioral fidelity. Future work should explore the influence of microbiome diversity on lifespan and stress resilience while refining methodologies for cricket rearing from egg to adulthood. By bridging invertebrate and vertebrate aging research, this study positions house crickets as a scalable, high-throughput model for investigating age-related functional decline, behavioral plasticity, and lifespan-extending interventions. Integrating behavioral assays, biomechanical analyses, and molecular markers of aging will further elucidate the interplay between morphology, function, and longevity, advancing the utility of crickets in comparative geroscience.

## Introduction

Aging is a universal process characterized by the progressive decline of physiological and morphological function, ultimately influencing an organism’s survival and fitness [1–2]. Understanding the patterns and mechanisms underlying age-related phenotypic deterioration is essential for elucidating the biology of aging and its implications for locomotion, sensory function, and overall health. While vertebrate models, particularly rodents, have been extensively studied in this regard [3–5], invertebrates remain comparatively understudied despite their potential to provide critical insights into the evolution and mechanisms of age-related functional decline [6–7]. Notably, few studies have systematically examined invertebrates with a direct translational relevance to vertebrate aging models, limiting their potential as complementary systems in biomedical research.

Aging phenotypes exhibit substantial variability within species due to genetic, environmental, and stochastic influences [8]. Documenting how key morphological traits shift across the lifespan is crucial for understanding age-related functional changes. Observable phenotypes such as body size, appendage length, and limb morphology are fundamental to locomotor performance, resource acquisition, and mating success, making them valuable indicators of aging-related functional decline [9]. However, the extent to which these morphological changes occur and whether they exhibit sex-specific patterns remain unresolved in many invertebrate species [10].

The house cricket (*Acheta domesticus*), with its relatively short lifespan, distinct and translational developmental stages (Figure 1), well-characterized physiological processes, and ease of maintenance in laboratory settings [11], serves as a practical model for studying morphological and functional deterioration associated with aging [12]. Its accessibility, well-defined biomechanics, and behavioral relevance further positions this insect as a promising yet underutilized model in aging research [13–14]. However, maximizing their translational potential requires standardizing husbandry protocols, as variability in housing conditions, diet composition, and population density can introduce confounding factors that obscure biological signals, ultimately limiting the reliability of experimental outcomes. This optimization is particularly crucial in aging research, where environmental factors significantly influence lifespan, behavior, and physiological decline. Despite their use in neurobiological and locomotor research, no studies have comprehensively examined morphological changes across the lifespan of crickets or assessed potential sex differences in these trajectories.

**Figure 1.**
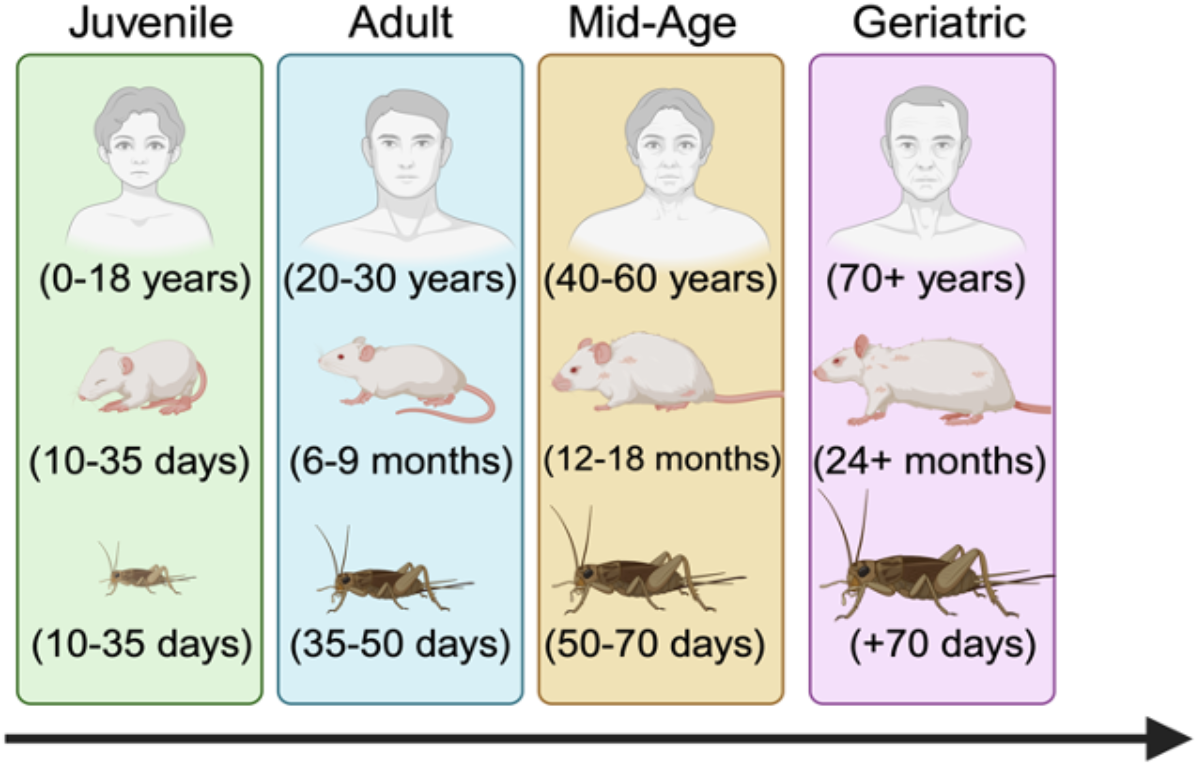
Translational relevance of the house cricket. Created in BioRender. Liao, G. (2025) https://BioRender.com/m17i385.

In the present study, we systematically quantify age-related changes in key morphological traits, including body weight, body length, antennal length, hind leg length, femoral cross-sectional area, femoral volume, and femoral surface area-to-volume ratio. Additionally, we assess proportional scaling relationships (e.g., hind leg length to body length, antennal length to body length, hind leg length to weight, body length to weight) and sensory to limb scaling ratios (e.g., hind leg to antennal length, antennal length to weight). By evaluating whether these traits exhibit significant age-related changes and sex-dependent patterns, this study provides novel insights into morphological aging in house crickets.

Furthermore, by integrating standardized husbandry practices, we enhance the reproducibility of cricket-based aging studies, reinforcing their potential as a translational model for aging and neurodegenerative research.

## Materials and Methods

### Genetic background & standardized handling techniques

House crickets with a heterogeneous genetic background were reliably sourced from Fluker Farms Inc. (Louisiana, USA), an approved commercial supplier which maintains a genetically diverse “American Mix” lineage bred over 60 generations from multiple regional strains across the United States. This genetic variability minimizes strain-specific artifacts, enhancing the model’s translational relevance for aging studies [15]. Furthermore, as these crickets are bred under controlled conditions from these vendors, this significantly reduces the risk of zoonotic disease transmission. While wild-caught crickets can harbor pathogens such as *E. coli*, as well as parasites like horsehair worms [16–17], commercially raised crickets undergo rigorous quality control measures to ensure they are free from such contaminants. Proper handling practices, including the use of gloves and maintaining clean enclosures, further mitigated any potential risks [18].

Standardized handling techniques are critical for minimizing stress and ensuring reproducibility in experimental paradigms. To ensure viability and experimental consistency, shipments were temperature-optimized and dispatched overnight only under suitable environmental conditions, reducing transport-induced stress [19]. Crickets were then safely transferred using modified culture tubes with holes drilled throughout the tube to provide proper ventilation (Figure 2) or small plastic containers covered at one end. These methods have been shown to reduce handling-related stress responses and physical injury in rodent models [20]. The tools used can also easily be sanitized and reduces the need for anesthesia for transport and routine weighing. For longitudinal studies requiring individual identification, researchers can employ a combination of sex-specific morphological differences (e.g., presence of wings in males, ovipositor in females) [21], distinct anatomical features (e.g., cerci and antennae length) [22], and weight measurements to track individual variation over time [23].

**Figure 2.**
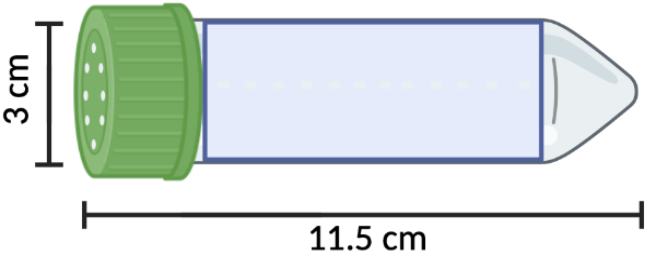
Standardized cricket handling technique using a modified test tube. Created in BioRender. Liao, G. (2025) https://BioRender.com/m17i385.

### Environmental conditions and housing

House crickets were reared under precisely controlled environmental conditions to optimize health, longevity, and behavioral stability. Temperature was maintained at 29 ± 1°C, with stable humidity levels (32 ± 3%) to mimic their natural habit and support normal physiological function [11,24]. Crickets were housed in a custom-designed double enclosure system designed to prevent escape while allowing for self-regulated photoperiods, which more accurately reflect their natural circadian rhythms [25]. The outer cage (80 cm x 60 cm x 60 cm) served to maintain stable temperature and humidity, while the inner enclosures (20 cm x 15 cm x 10 cm) allowed for controlled light exposure and behavioral monitoring. For cleaning feces and scattered food, the bottoms of the cages were scraped once a week to avoid causing significant anxiety to the crickets [26].

To provide environmental enrichment and prevent overcrowding stress, egg cartons were placed within the cages, offering shaded areas that crickets could autonomously select based on their preferred microenvironment (Figure 3). This self-determined photoperiod structure has been shown to stabilize behavioral patterns and self-regulated circadian rhythms, improve physiological health, and enhance reproductive success compared to standardized light cycles, which can disrupt activity patterns and stress adaptation [27]. Housing densities were maintained at 20–30 individuals per cage to optimize social interactions while minimizing aggressive behaviors associated with overcrowding and resource competition [28–29]. Both sexes were co-housed to reflect natural environmental conditions, which promotes normal social and mating behaviors [30].

**Figure 3.**
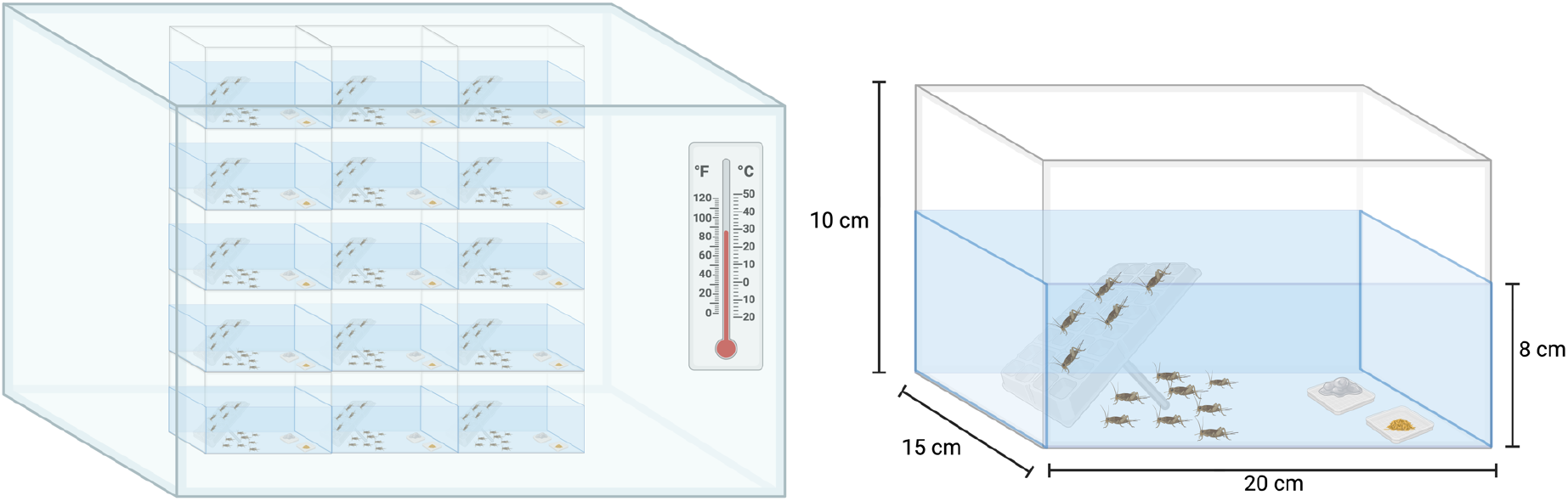
Schematic representation of the double enclosure system for housing crickets (left). Detailed dimensions and design features of the inner cricket habitat are provided (right). Created in BioRender. Liao, G. (2025) https://BioRender.com/m17i385.

Oviposition substrate was deliberately excluded to prevent reproductive variability from confounding our findings, as we are primarily focused on aging rather than reproductive investment. While oviposition sites can be present in natural environments, their availability is inconsistent, and previous studies suggest that their absence may influence food intake and energy allocation in certain cricket morphs [31]. By standardizing environment conditions without an oviposition substrate, we aimed to minimize uncontrolled reproductive effects and ensure that observed differences were attributable to aging rather than reproductive status.

Unlike traditional laboratory animal models maintained in specific pathogen-free (SPF) conditions, crickets were housed under conventional conditions to preserve natural microbial exposures, thereby increasing translational relevance [32]. Research in mammalian models has demonstrated that exposure to environmental pathogens significantly alters immune function, rendering laboratory animals more physiologically comparable to wild populations and, by extension, human subjects [33].

### Diet and nutrition

A standardized diet was provided to all crickets to ensure consistent nutritional intake and minimize variability in metabolic and physiological outcomes. The diet was formulated by combining Picolab Rodent Diet 20 (5053, Irradiated; Purina Mills, USA, which contains corn, soybean, wheat, fish meal, and a vitamin-mineral mixture), with gelatin as a binding agent. The control diet was prepared by dissolving gelatin in water at 100°C, cooling to approximately 60°C, and subsequently mixing it with rodent chow. This homogenized mixture was then refrigerated at 20°C overnight before undergoing dehydration in a standard food dehydrator to deter mold growth and extend shelf life. Once dried, the diet was further homogenized using a food processor to maintain consistency and prevent selective feeding behavior.

Crickets were provided *ad libitum* access to both food and water, with hydration supplied by water gel packs (Napa Nectar Plus, Systems Engineering Lab Group, Napa, CA), a gel-based medium that minimizes spillage and accidental drowning while maintaining stable humidity levels within enclosures. The water gels typically remained viable for 3-4 days before requiring replacement due to dehydration or contamination. Similarly, food replacement frequency varied depending on cricket density and age; for example, a cohort of 30 six-week-old crickets typically required 2 g of food replenishment every 3-4 days, whereas smaller or older groups necessitated replacement every 5-6 days (unpublished observations).

Cricket survival was monitored daily, with deceased individuals promptly removed to prevent contamination and discourage cannibalism.

### Age classification and selection criteria

Crickets were classified into four distinct age groups based on post-hatch developmental stages: juveniles (4 weeks old), adults (6 weeks old, post-molt cessation), mid-aged (8 weeks old), and geriatric (≥10 weeks old). To account for potential genetic and environmental variability, individuals were selected from multiple populations across independent studies conducted within the laboratory. Only morphologically intact individuals without visible injuries or deformities were included in downstream analyses.

### Morphometric measurements

Live crickets were individually placed in a covered jar and weighed to the nearest 0.001 g using a precision analytical balance (Mettler Toledo XPR205, Switzerland). Following euthanasia, a series of morphometric measurements were conducted to quantify age-related phenotypic changes. Body length was measured using flat acrylic rulers, excluding cerci, wings (males only), and ovipositor (females only) to ensure standardized comparisons. Both antennae were measured, and in cases of asymmetry, an average length was recorded. Hind leg length was assessed by fully extending the leg parallel to the measuring surface, mitigating potential bias introduced by post-mortem joint rigidity.

### Femur morphometric analysis and structural modeling

To investigate biomechanical and structural adaptations in the hind leg femur across developmental stages, femoral morphology was quantified using a hybrid geometric model approximating the femur as an ellipsoidal core with a truncated ellipsoidal conical end (Figure 4). This model optimally balances computational accuracy with biological realism, allowing for high-fidelity representation of cricket femoral architecture. Morphometric calculations included femoral volume, cross-sectional area (CSA), and surface area-to-volume (SA/V) ratio, as detailed below.

**Figure 4.**
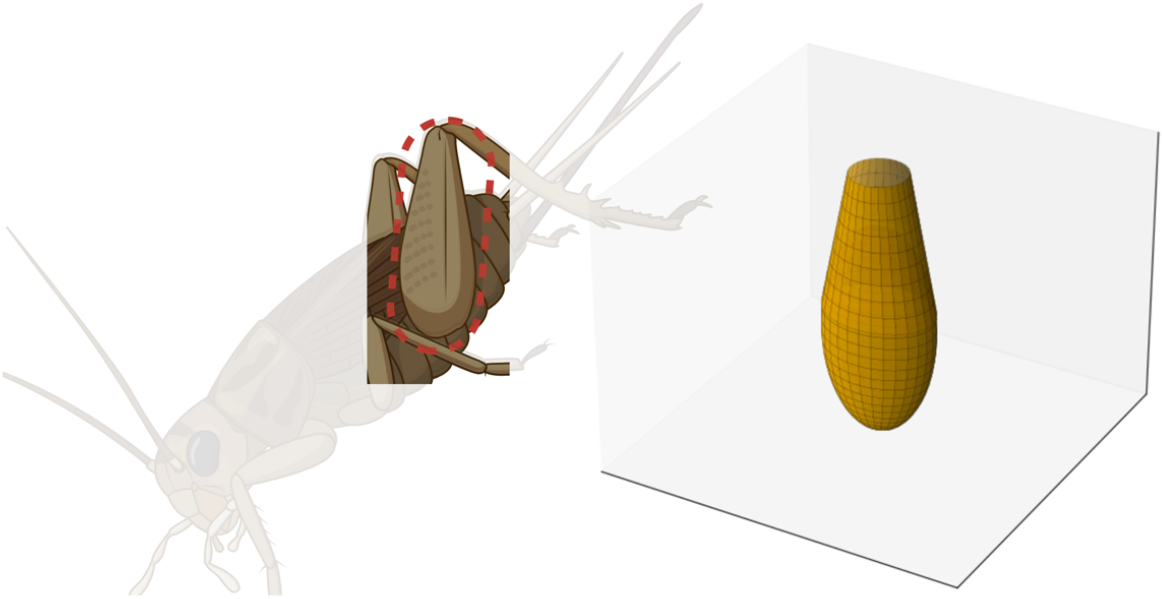
Schematic and geometric hybrid model of the cricket hind leg femur. Comparison of the schematic diagram of a cricket’s hind leg femur, highlighted with a red dotted circle (left) and the corresponding geometric hybrid model (right). The schematic was created in BioRender. Liao, G. (2025) https://BioRender.com/m17i385, while the 3D model was generated using Python 3.13.2 with Matplotlib.

### Femur Volume Calculation

To estimate the femur volume (*V*), we modeled the structure as a composite geometric shape consisting of a proximal ellipsoid and a distal truncated ellipsoidal cone. To simplify calculations, we defined the truncation point at the midpoint of the femur, where the radius of the ellipsoid and the truncated cone are equal. Thus, the total volume is given by:

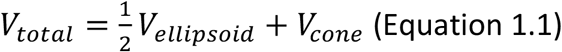

The volume of the ellipsoid was calculated using the standard ellipsoid volume formula:

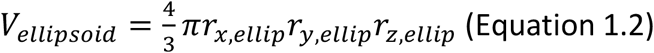

For the truncated ellipsoidal cone, we assume that each cross-section perpendicular to the femur’s longitudinal axis is an ellipse, with the major and minor axis lengths varying linearly along the femur’s height. The volume was obtained by integrating the product of the semi-major and semi-minor axis functions across the height of the truncated cone:

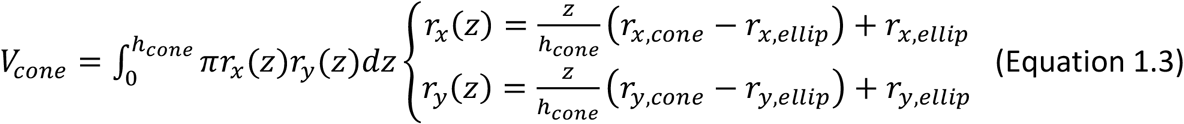

Solving the integral yields the final volume expression:

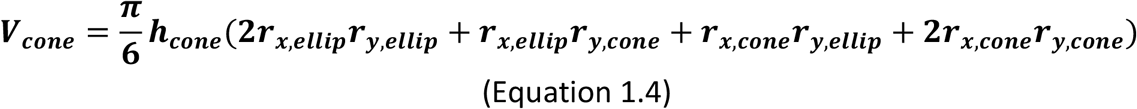

### Femur Cross-Sectional Area Calculation

To estimate the largest cross-sectional area (*A*(*x*)) of the femur along its length, we assumed an elliptical cross-section, similar to the volume model. The general equation for the area of an ellipse is:

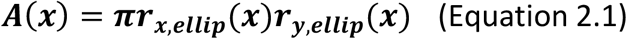

### Femur Surface Area Calculation

The femur surface area (SA) was estimated by summing the contributions from the proximal ellipsoid and the distal truncated ellipsoidal cone:

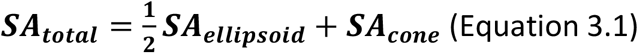

The surface area of the ellipsoid was computed using the Knud Thomsen approximation:

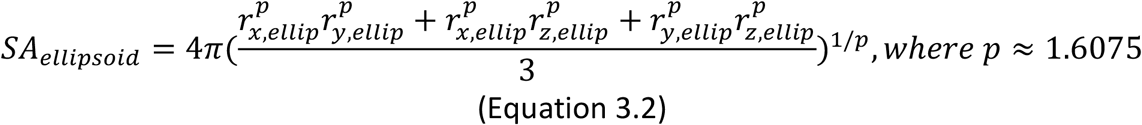

The surface area of the truncated ellipsoidal cone was calculated as follows:

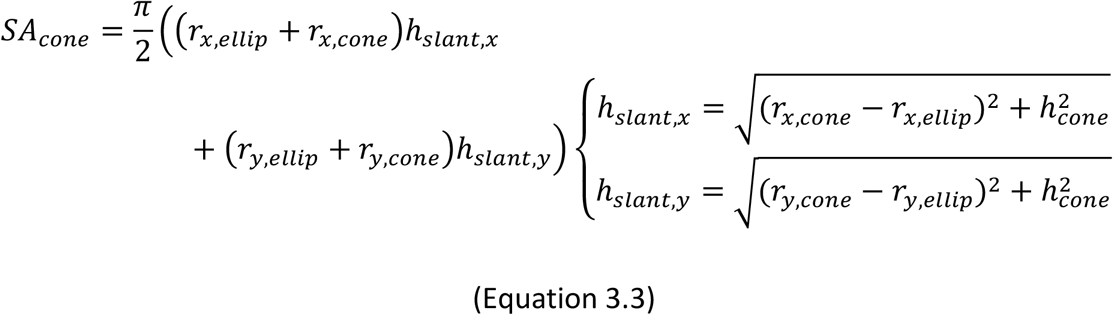

### Euthanasia and Sample Preparation

To ensure rapid and humane euthanasia, crickets were exposed to CO_2_ in a controlled chamber and subsequently decapitated as an adjunctive method following American Veterinary Medical Association (AVMA) guidelines [34]. Post-mortem assessments were conducted immediately after euthanasia to prevent degradation-related artifacts in morphometric analyses.

### Statistical Analysis

All data were stratified by age group and sex. Normality was assessed using the Shapiro-Wilk test, except in cases where the sample size was sufficiently large (N ≥ 30) to assume normality under the central limit theorem. Normally distributed data were analyzed using parametric tests, while non-normally distributed data were assessed using non-parametric methods. A one-way analysis of variance (ANOVA) was used to evaluate the independent effects of age and sex, with Tukey’s post-hoc test for multiple comparisons. A two-way ANOVA was performed to examine both main and interactive effects of age and sex. In cases where no significant interaction was observed but at least one main effect was significant, post-hoc comparisons were performed using Bonferroni-adjusted multiple comparisons for sex differences within each age group and Tukey’s multiple comparison test for differences between age groups within each sex. All statistical analyses were conducted using GraphPad Prism (version 10.0.3), with statistical significance set at α = 0.05.

## Results

### Body weight increases with age and is generally higher in females

Across age groups, body weight exhibited differences, with geriatric crickets weighing more than both adults and juveniles but not differing from mid-aged cohorts (Figure 5A). Mid-aged crickets were heavier than both adults and juveniles, while adults weighed more than juveniles. Sex-based comparisons revealed that females generally had higher body weights than males across all groups (Figure 5B). When stratified by sex and age, female crickets followed the same trend as the overall group (Figure 5C). However, in males, geriatric individuals were only heavier than juveniles and did not differ from adult or mid-aged crickets. In both sexes, mid-aged individuals maintained their weight advantage over younger groups, mirroring the overall trend. Within each age group, females were heavier than males in the adult, mid-age, and geriatric groups, whereas no sex differences were observed among juveniles (Figure 5D).

**Figure 5.**
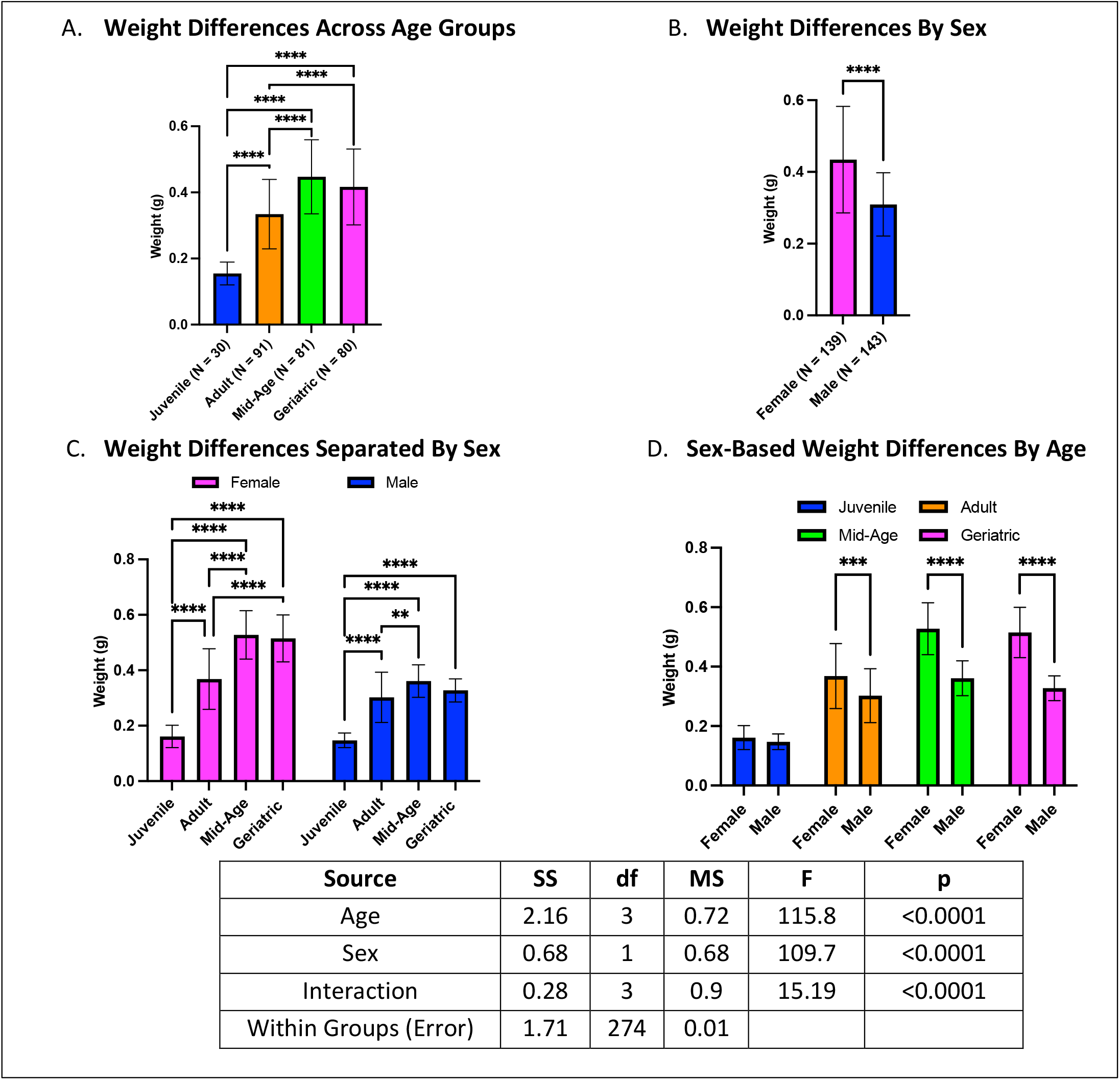
Age- and sex-related differences in body weight. **(A)** Geriatric crickets were heavier than juveniles and adults but not mid-aged crickets. **(B)** Females were generally heavier than males. **(C)** The weight trend remained consistent for females, while in males, geriatrics only differed from juveniles. **(D)** Sex-based differences were evident in adults, mid-aged, and geriatric crickets but not in juveniles (**p < 0.01, ***p < 0.001, ****p < 0.0001).

### Age-related changes in antennal length are independent of sex

Antennal length varied with age but showed no overall sex-based differences (Figure 6A-B). Geriatric crickets had longer antennae than mid-aged and juvenile crickets but did not differ from adults (Figure 6A). Mid-aged crickets had shorter antennae than adults but longer than juveniles, while adults had longer antennae than juveniles. When stratified by sex, females followed the same age-dependent pattern as the overall analysis (Figure 6C). For males, however, geriatric and mid-aged individuals did not differ, nor did mid-aged and adult crickets. Sex comparisons within each age group revealed no differences (Figure 6D).

**Figure 6.**
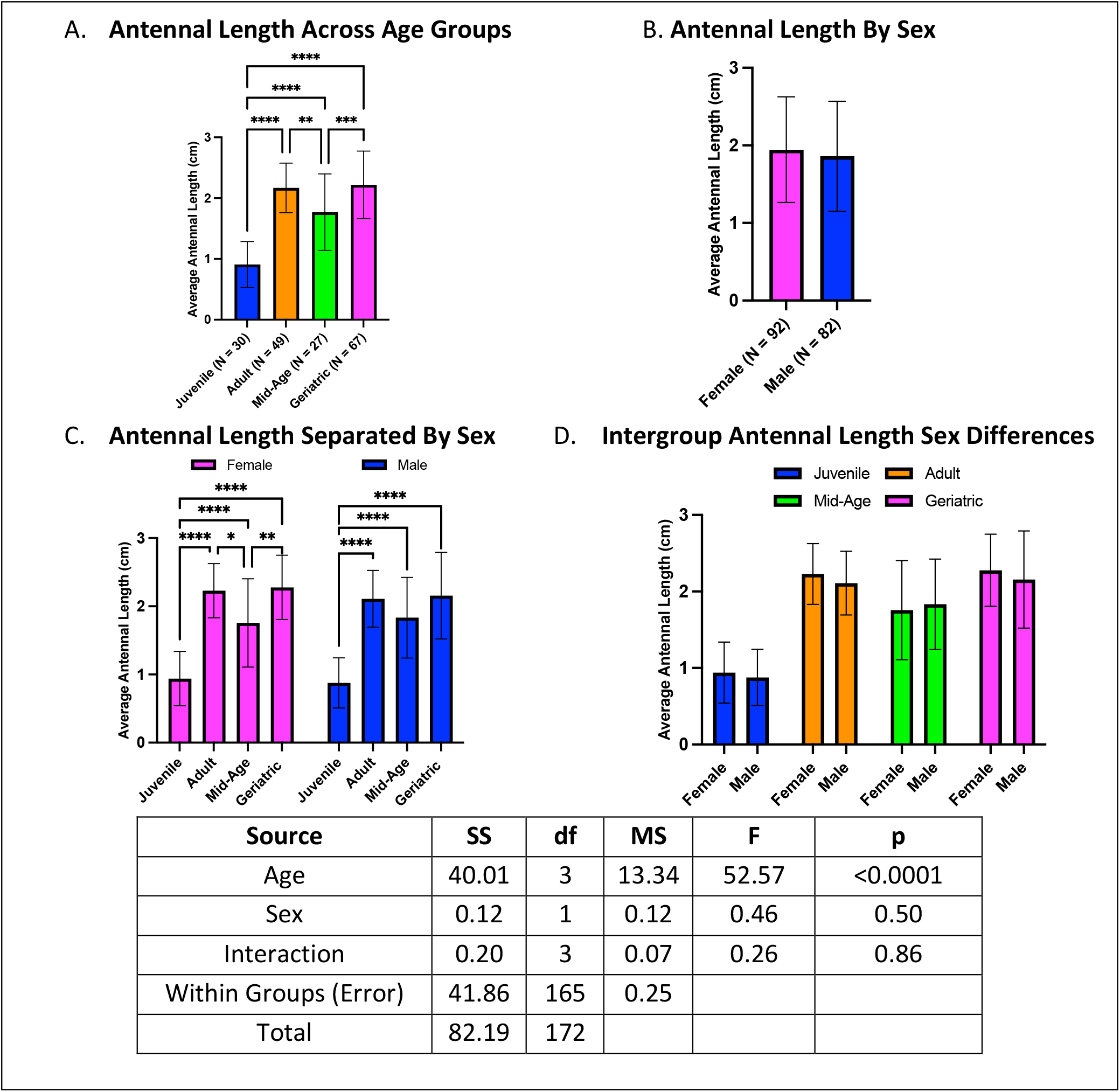
Age-and sex-related differences in antennal length. **(A)** Geriatric crickets had longer antennae than juveniles and mid-aged crickets but not adults. **(B)** No overall sex differences were observed. **(C)** While females followed the same trend as the overall group, males showed no difference between geriatric and mid-aged individuals. **(D)** No sex differences were detected within age groups (*p < 0.05, **p < 0.01, ***p < 0.001, ****p < 0.0001).

### Body length increases with age and is greater in females

Body length increased with age, with geriatric crickets being larger than both adults and juveniles, but not different from mid-aged crickets (Figure 7A). Similarly, mid-aged crickets were longer than adults and juveniles, and adults were longer than juveniles. Females generally had longer bodies than males (Figure 7B). When analyzing sex differences by age, both male and female crickets followed the same trends observed in the overall comparisons (Figure 7C). Within individual age groups, sex-based differences emerged in mid-aged and geriatric crickets, with females being longer than males. However, no differences were detected in juvenile or adult cohorts (Figure 7D).

**Figure 7.**
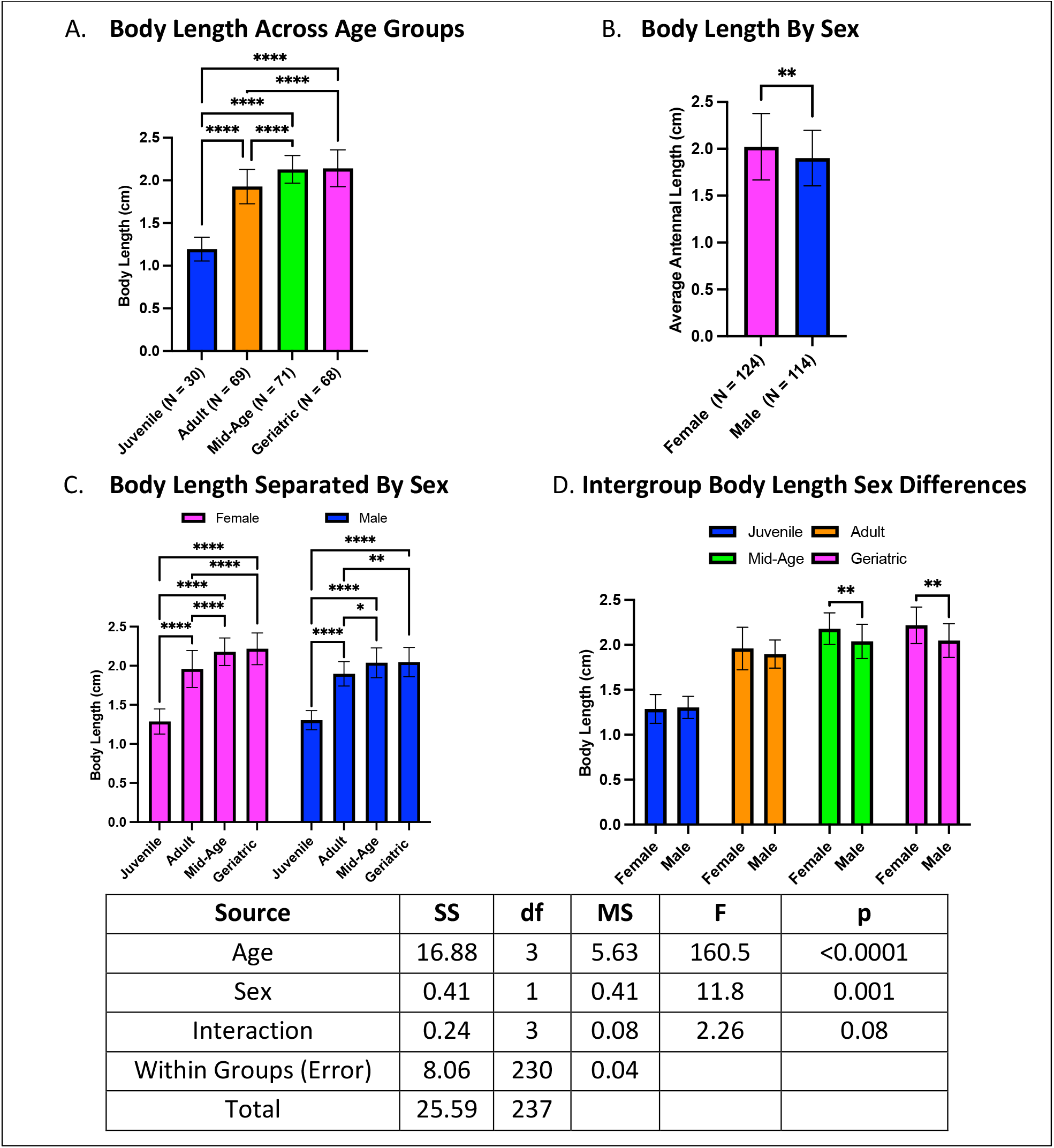
Age- and sex-related differences in body length. **(A)** Geriatric and mid-aged crickets had longer bodies than younger groups. **(B)** Females generally had longer body lengths than males. **(C)** Both males and females followed the same age-related trends. **(D)** Sex differences were observed in mid-aged and geriatric groups but not in juveniles or adults (*p < 0.05, **p < 0.01, ****p < 0.0001).

### Hind leg length is modulated by age and sex

Hind leg length exhibited distinct age-related trends, with geriatric crickets having longer hind legs than juveniles but not differing from mid-aged or adult crickets (Figure 8A). Mid-aged crickets had longer hind legs than both adults and juveniles, while adults had longer hind legs than juveniles. Females generally had longer hind legs than males across all groups (Figure 8B). Within sex-stratified comparisons, geriatric females had shorter hind legs than mid-aged females but longer than juveniles, with no difference from adults (Figure 8C). Mid-aged females had longer legs than both adults and juveniles, while adult females had longer legs than juveniles. Male crickets followed a similar trend, except that geriatric males did not differ from mid-aged males. Sex comparisons within each age group revealed no differences in hind-leg length among juveniles or geriatric crickets (Figure 8D). However, in mid-aged and adult groups, females had longer hind legs than males.

**Figure 8.**
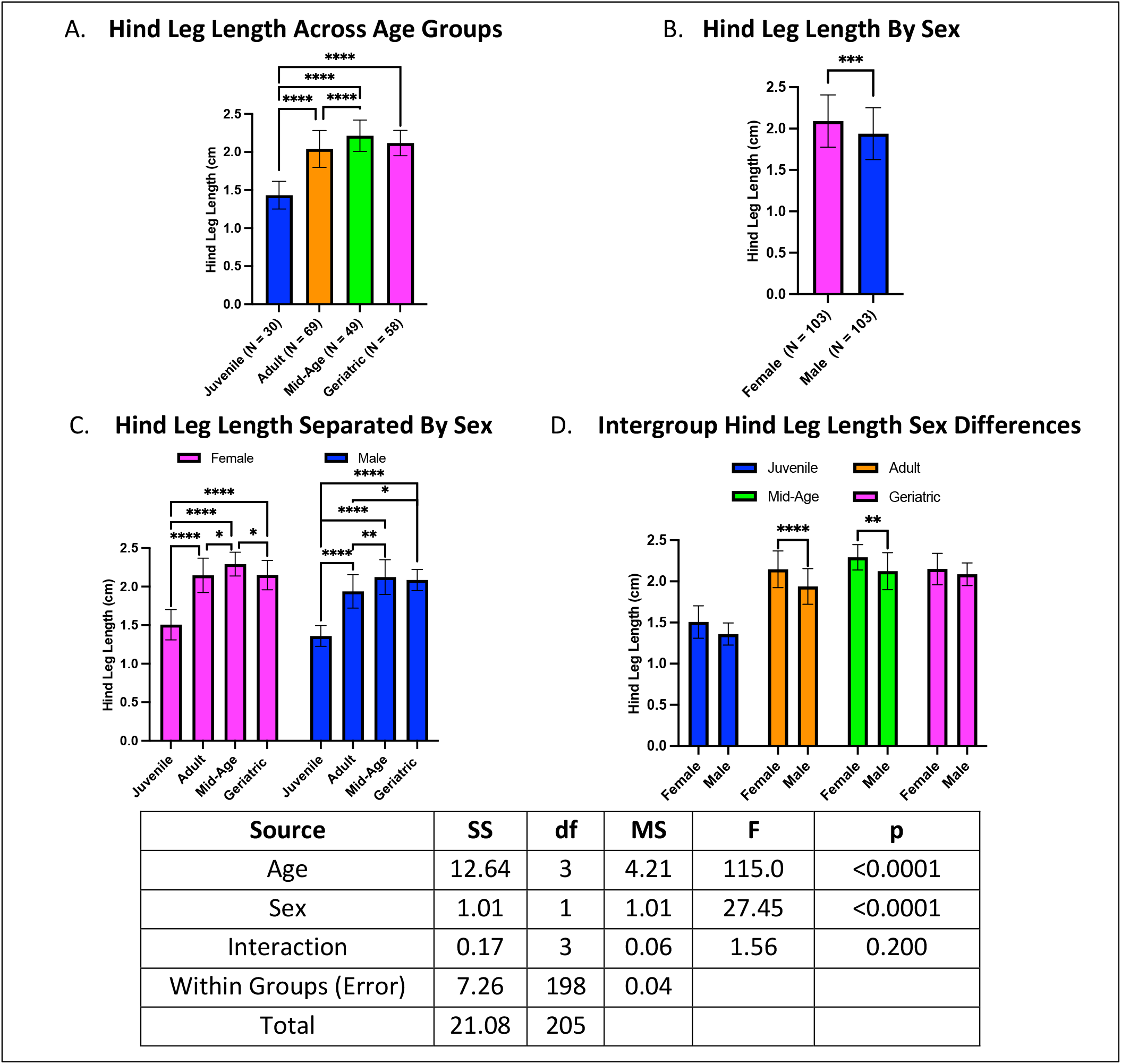
Age- and sex-related differences in hind leg length. **(A)** Geriatric crickets had longer hind legs than juveniles but did not differ from mid-aged or adult crickets. **(B)** Females generally had longer hind legs than males. **(C)** The geriatric and mid-aged females exhibited distinct trends, with mid-aged females having the longest hind legs. **(D)** Sex-based differences were evident in mid-aged and adult groups but not in juveniles or geriatrics (*p < 0.05, **p < 0.01, ***p < 0.001, ****p < 0.0001).

### Femoral volume increases with age, with sex-specific differences emerging in adulthood

Femoral volume exhibited age-related changes, with geriatric crickets displaying larger femoral volumes than juveniles and adults, while mid-age crickets had the largest femoral volumes overall (Figure 9A). Adults also had larger femoral volumes than juveniles. When analyzed by sex, females consistently exhibited larger femoral volumes than males (Figure 9B). Further stratification by both age and sex revealed that both sexes followed the same trend as the overall data, except mid-age males did not differ from adults (Figure 9C). Within-group sex differences were absent in juvenile and adult crickets, but among mid-age and geriatric crickets, females exhibited larger femoral volumes than their male counterparts (Figure 9D). These findings suggest that femoral volume increases with age, with the most pronounced differences occurring during mid-age, and that sexual dimorphism in femoral volume emerges after the juvenile stage.

**Figure 9.**
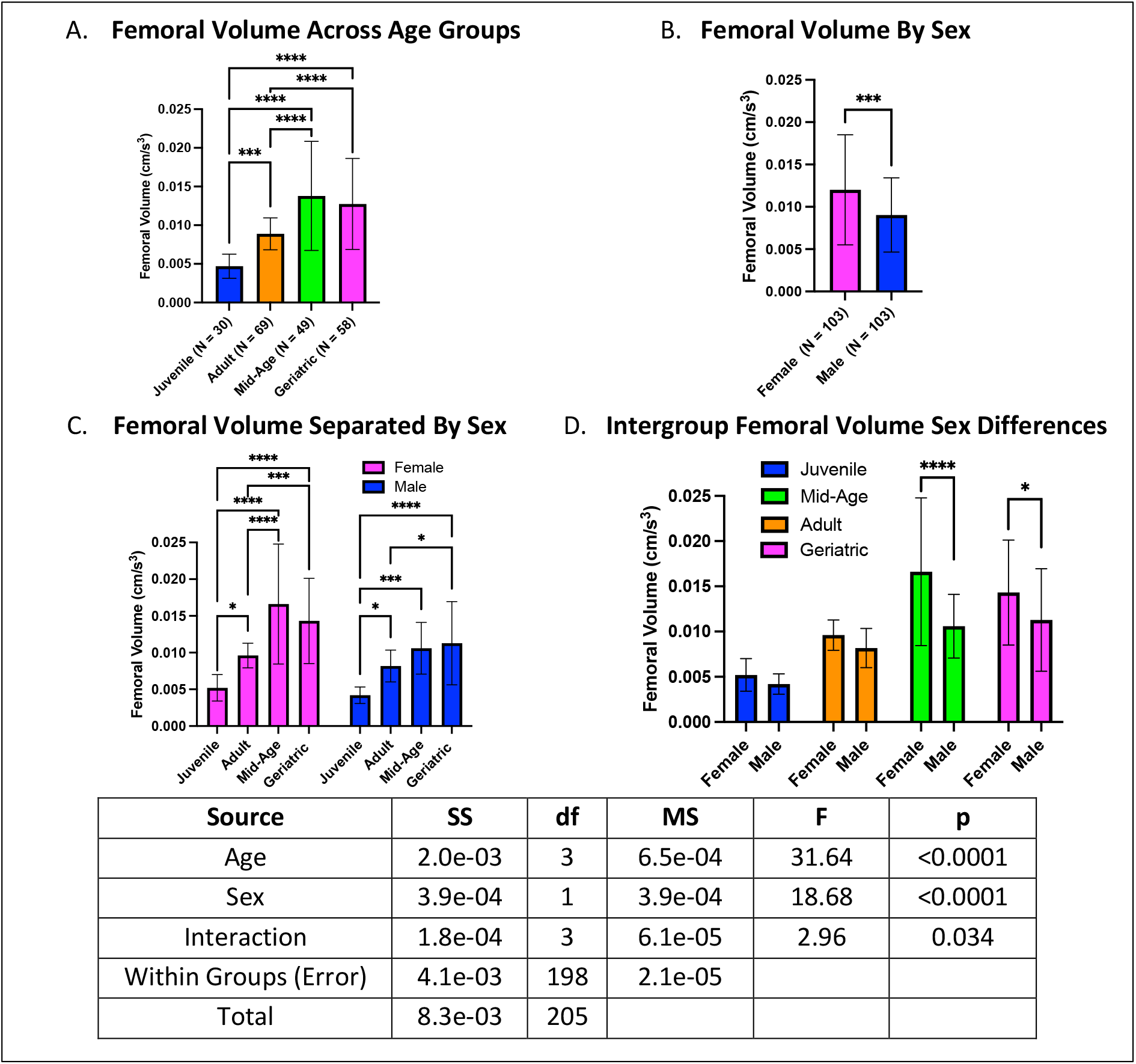
Age- and sex-dependent differences in femoral volume. **(A)** Mid-age crickets exhibited the largest femoral volume, followed by geriatric and adult crickets, with juveniles having the smallest femoral volumes. **(B)** Females generally had larger femoral volumes than males. **(C)** Mid-age crickets had larger femoral volumes than adults and juveniles, while adults exhibited larger femoral volumes than juveniles. **(D)** Females exhibited larger femoral volumes than males in mid-age and geriatric groups (*p < 0.05, ***p < 0.001, ****p < 0.0001).

### Femoral cross-sectional area (CSA) increases with age, with no intra-sex differences

Femoral CSA exhibited age-related increases, with geriatric crickets displaying larger CSA than juveniles and adults but showing no difference from mid-age crickets. Mid-age crickets had femoral CSAs that surpassed both adults and juveniles, while adults exhibited larger CSA than juveniles (Figure 10A). Overall, females had larger femoral CSA than males (Figure 10B). When stratified by both age and sex, the patterns remained consistent with the overall analysis, except that for females, juvenile and adult crickets no longer differed while for males, adults no longer differed from mid-age or geriatric crickets (Figure 10C). Within-group sex comparisons showed females had larger femoral CSA than males in mid-age cohorts only (Figure 10D).

**Figure 10.**
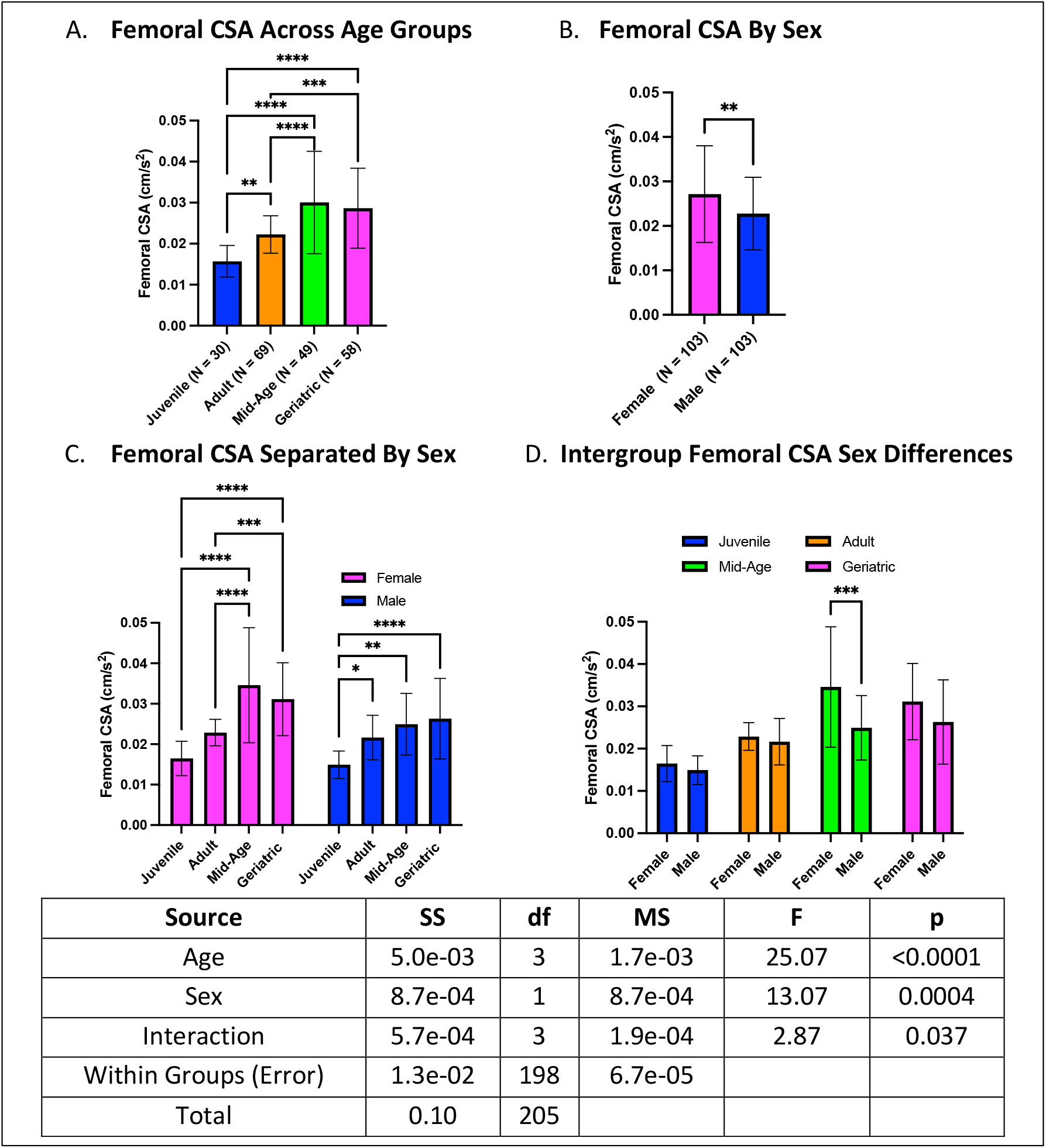
Age- and sex-dependent differences in femoral CSA. **(A)** Mid-age crickets exhibited the largest CSA, followed by geriatric and adult crickets, with juveniles having the smallest CSA. **(B)** Females generally had larger CSA than males. **(C)** Compared to overall differences, females and males showed distinct age-related changes. **(D)** Mid-age females had larger femoral CSA than males (*p < 0.05, **p < 0.01, ****p < 0.0001).

### Femoral surface area-to-volume ratio (SA/V) decreases with age, with males exhibiting higher ratios than females

Femoral SA/V ratio showed an inverse relationship with age, with juveniles having higher SA/V ratios than all other age groups. Adults had higher SA/V ratios than mid-age and geriatric crickets, while geriatric and mid-age crickets did not differ (Figure 11A). Males had consistently higher SA/V ratios than females (Figure 11B). When analyzed by both age and sex, females followed the same trend as overall analysis while for males, juveniles maintained higher SA/V ratios than all other groups without any additional differences observed between mid-age, adult, and geriatric crickets (Figure 11C). Within-group sex comparisons showed mid-age and geriatric males maintained higher SA/V ratios than females (Figure 11D).

**Figure 11.**
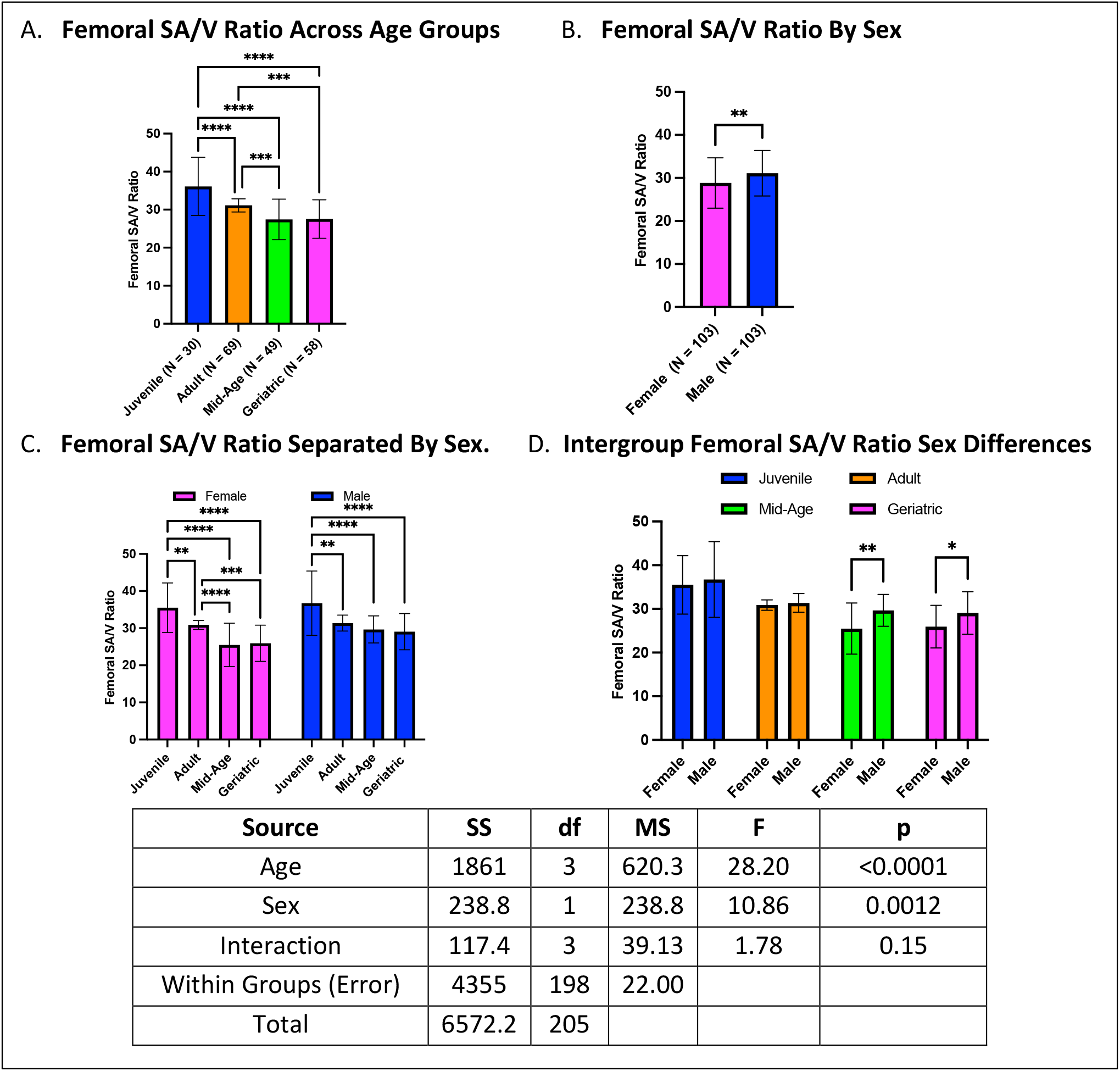
Age- and sex-dependent differences in femoral SA/V ratio. **(A)** Juveniles exhibited the highest SA/V ratios, followed by adults, mid-age, and geriatric crickets. **(B)** Males generally had higher SA/V ratios than females. **(C)** Females showed the same trend as overall analysis while for males, juveniles maintained higher SA/V ratios, while no other age groups differed. **(D)** Mid-age and geriatric males had higher SA/V ratios than females (*p < 0.05, **p < 0.01, ***p < 0.001, ****p < 0.0001).

### Hind leg to body length ratio declines with age, particularly in females

Hind leg to body length ratios showed a decline in geriatric groups compared to both adults and juveniles but did not differ from mid-age crickets (Figure 12A). No differences were observed between sexes in the overall analysis (Figure 12B). When stratified by both age and sex, males exhibited no differences across age groups, whereas females demonstrated a pronounced decline in hind leg-to-body length ratio with aging. Specifically, geriatric females had lower ratios than adults and juveniles but did not differ from mid-age crickets. Additionally, mid-age females had lower ratios than juveniles (Figure 12C). Within-group sex comparisons revealed that in younger cohorts (juveniles and adults), females exhibited larger ratios than males. However, this sex difference was not observed in the older cohorts (Figure 12D).

**Figure 12.**
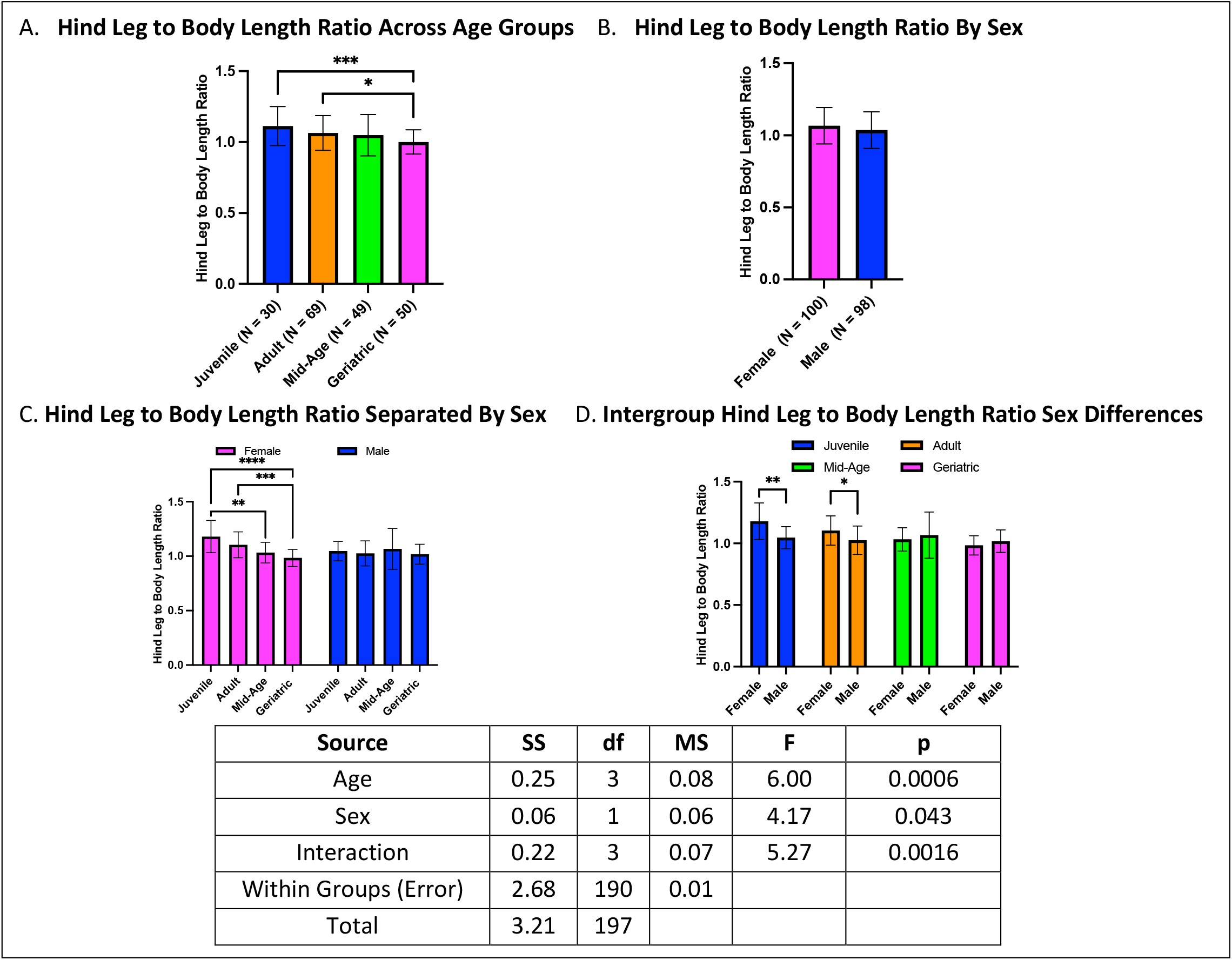
Hind leg to body length ratios across age and sex. **(A)** Geriatric crickets had lower hind leg-to-body length ratios compared to juveniles and adults but did not differ from mid-age crickets. **(B)** No sex differences were observed in overall comparisons. **(C)** When stratified by both age and sex, females but not males showed age-related declines. **(D)** Sex differences were apparent in juveniles and adults only, with females exhibiting higher ratios than males (*p < 0.05, **p < 0.01, ***p < 0.001, ****p < 0.0001).

### Antenna to body length ratio increases with age in both sexes

Antenna to body length ratios exhibited an increase in geriatric crickets compared to both mid-age and juvenile cohorts. Adults also had higher ratios than mid-age and juvenile crickets (Figure 13A). No sex differences were detected in overall comparisons (Figure 13B). When analyzed by both age and sex, females followed a similar trend as the overall analysis, with juveniles exhibiting lower ratios than adults and geriatrics. However, mid-age females had lower ratios than adults but did not differ from geriatrics. In contrast, males only showed differences between juveniles and the older cohorts, with adults and geriatrics having similarly high ratios (Figure 13C). Within-group sex comparisons revealed no differences in any age group (Figure 13D).

**Figure 13.**
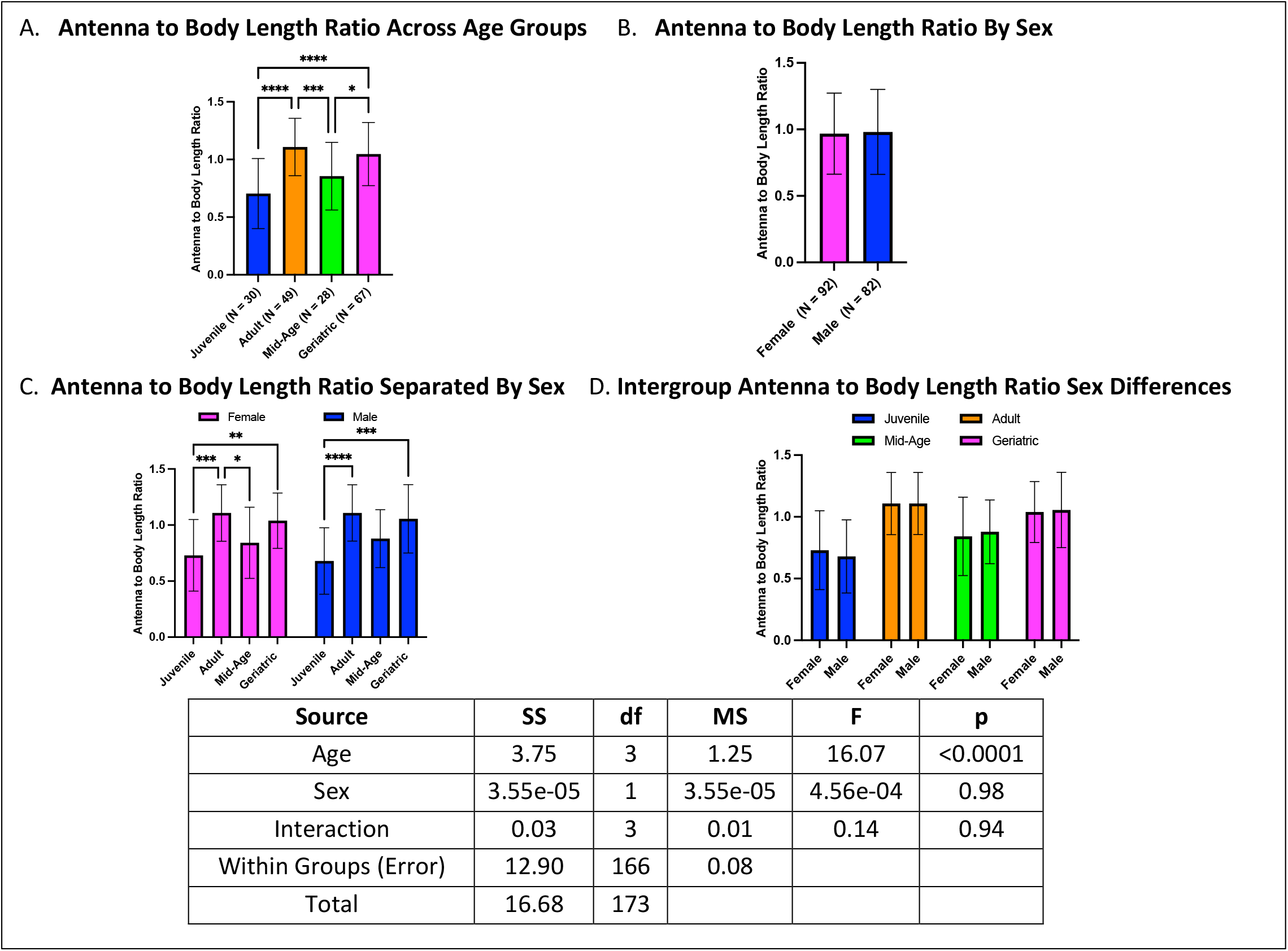
Antenna to body length ratios across age and sex. **(A)** Geriatric crickets had higher antenna to body length ratios than juveniles and mid-age crickets, with adults also showing elevated ratios. **(B)** No sex differences were seen in overall analysis. **(C)** Females exhibited a progressive increase with age, while males only had differences between juveniles and the older cohorts. **(D)** No sex differences were observed within individual age groups (*p < 0.05, **p < 0.01, ***p < 0.001, ****p < 0.0001).

### Hind leg length to weight decreases with age, with males exhibiting higher ratios

Hind leg length to weight ratios declined with age, with mid-age and geriatric crickets exhibiting the lowest ratios, both lower than juveniles and adults. Additionally, adults had lower ratios than juveniles (Figure 14A). A sex effect was observed, with males exhibiting higher hindleg-to-weight ratios than females (Figure 14B). When further stratified by age and sex, both males and females followed the same trend as the overall analysis, with hind leg length to weight ratio decreasing progressively with age (Figure 14C). Within-group sex comparisons revealed that males had higher ratios than females in adults, mid-age, and geriatric cohorts, whereas no sex differences were observed in juveniles (Figure 14D).

**Figure 14.**
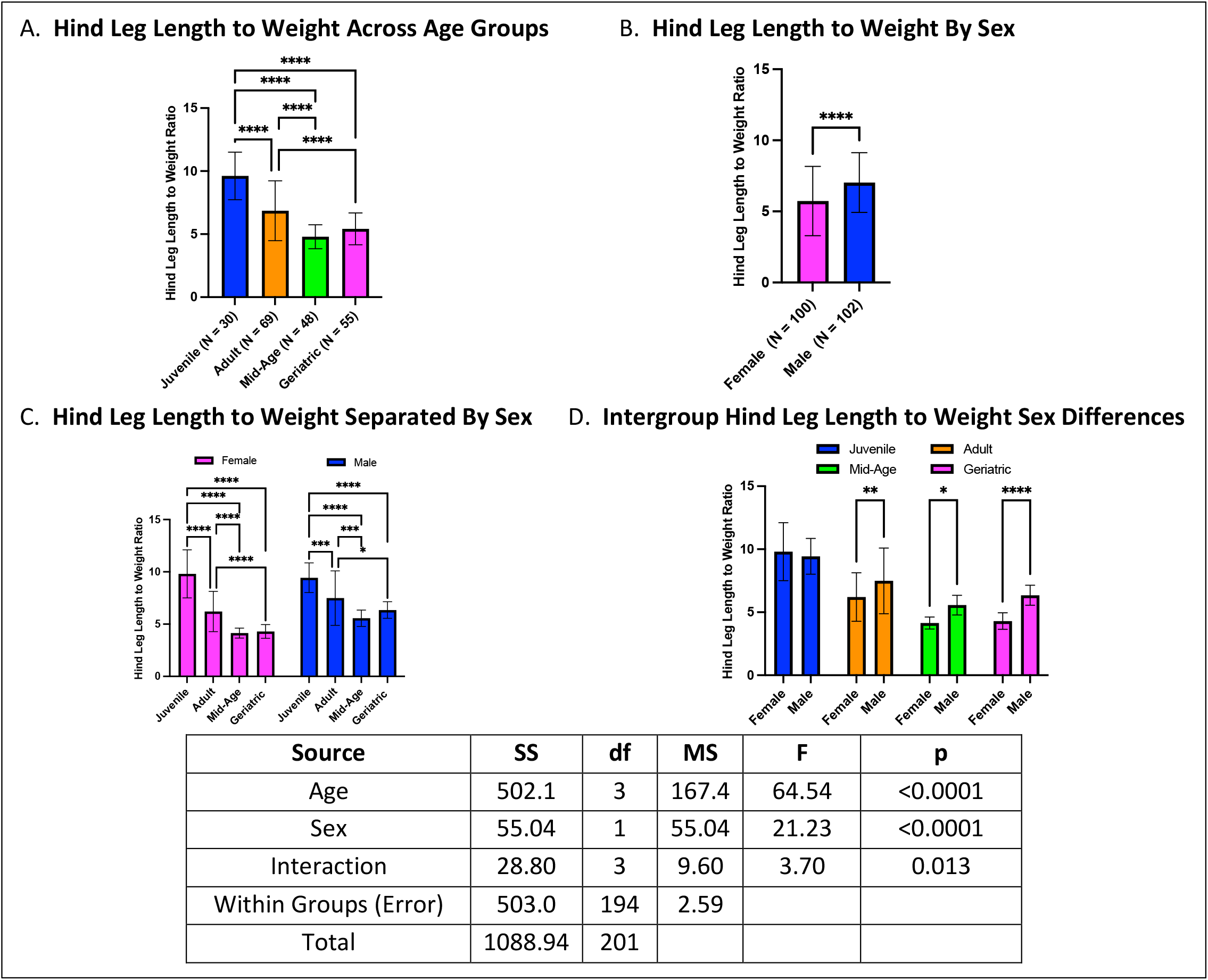
Hind leg length to weight ratios across age and sex. **(A)** Older cohorts (mid-age and geriatrics) exhibited lower hind leg length to weight ratios compared to juveniles and adults. **(B)** Males had overall higher hind leg length to weight ratios than females. **(C)** The same age-related decline is observed when analyzed separately by sex. **(D)** Sex differences are present in adults, mid-age, and geriatric crickets, but not in juveniles (*p < 0.05, **p < 0.01, ***p < 0.001, ****p < 0.0001).

### Body length to weight ratio decreases with age, with males exhibiting higher ratios

Body length to weight ratios declined with age, with both mid-age and geriatric crickets exhibiting lower ratios than adults and juveniles (Figure 15A). A sex difference was observed, with males having higher body length to weight ratios than females in overall comparisons (Figure 15B). When further stratified by age and sex, both males and females followed the same pattern as the overall analysis, with body length to weight ratios decreasing with age (Figure 15C). Within-group sex comparisons revealed that males had higher ratios than females in adult, mid-age, and geriatric cohorts, whereas no sex differences were observed in juveniles (Figure 15D).

**Figure 15.**
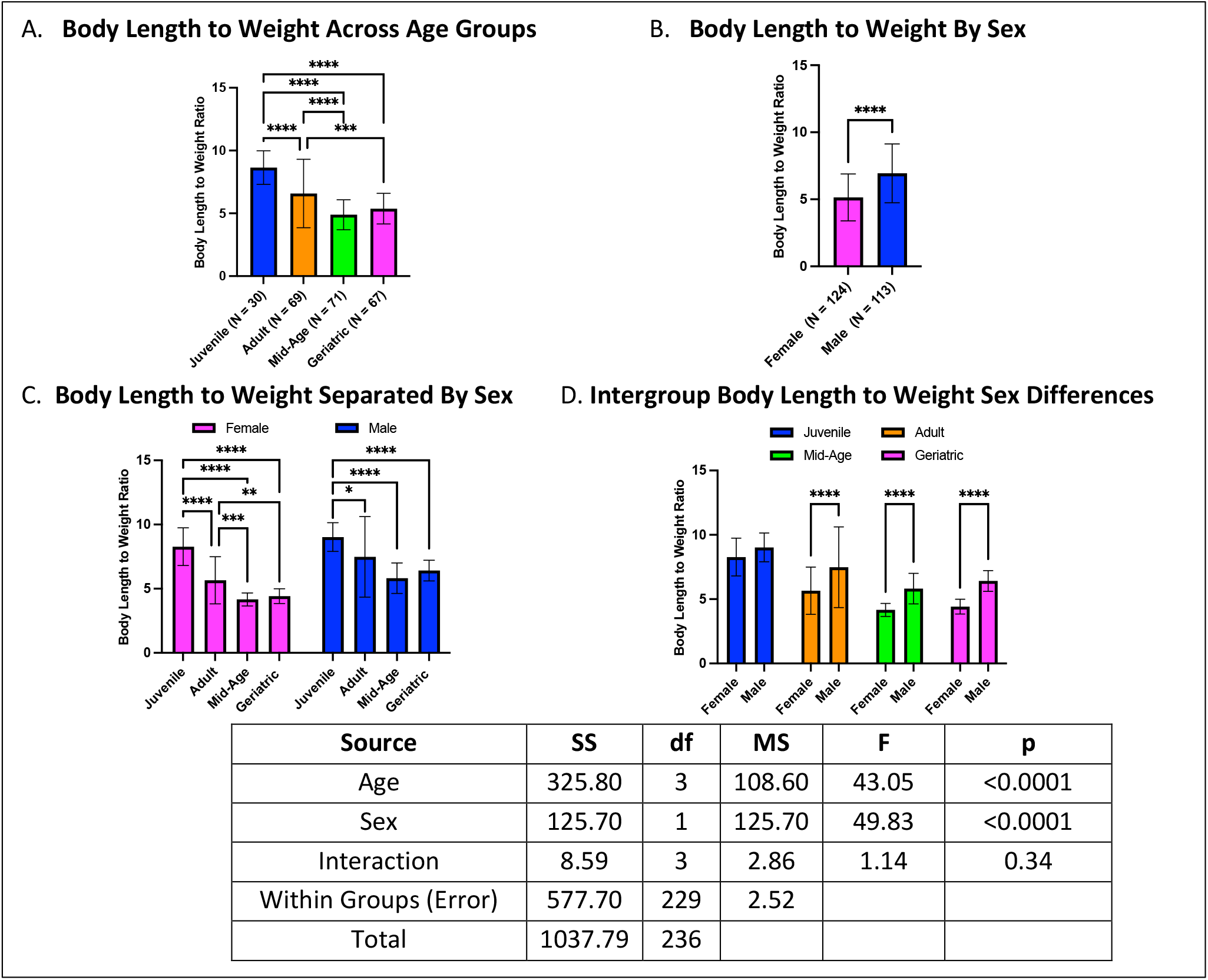
Body length to weight ratios across age and sex. **(A)** Mid-age and geriatric crickets exhibited lower body length to weight ratios compared to juveniles and adults. **(B)** Males had higher body length to weight ratios than females in overall comparisons. **(C)** When stratified by both age and sex, both males and females exhibited an age-related decline in body length to weight ratio. **(D)** Sex differences are present in adults, mid-age, and geriatric crickets, but not in juveniles (*p < 0.05, **p < 0.01, ***p < 0.001, ****p < 0.0001).

### Hind leg to antennal length ratio declines in adult and geriatric groups

Hind leg to antennal length ratios were lowest in adult and geriatric crickets, with both groups exhibiting lower ratios than mid-age and juvenile cohorts (Figure 16A). No sex differences were observed in the overall analysis (Figure 16B). When further stratified by age and sex, both males and females followed the same trend as the overall analysis, with juveniles exhibiting higher ratios than both adults and geriatrics but not differing from mid-age crickets (Figure 16C). Within-group sex comparisons revealed no differences across any age group (Figure 16D).

**Figure 16.**
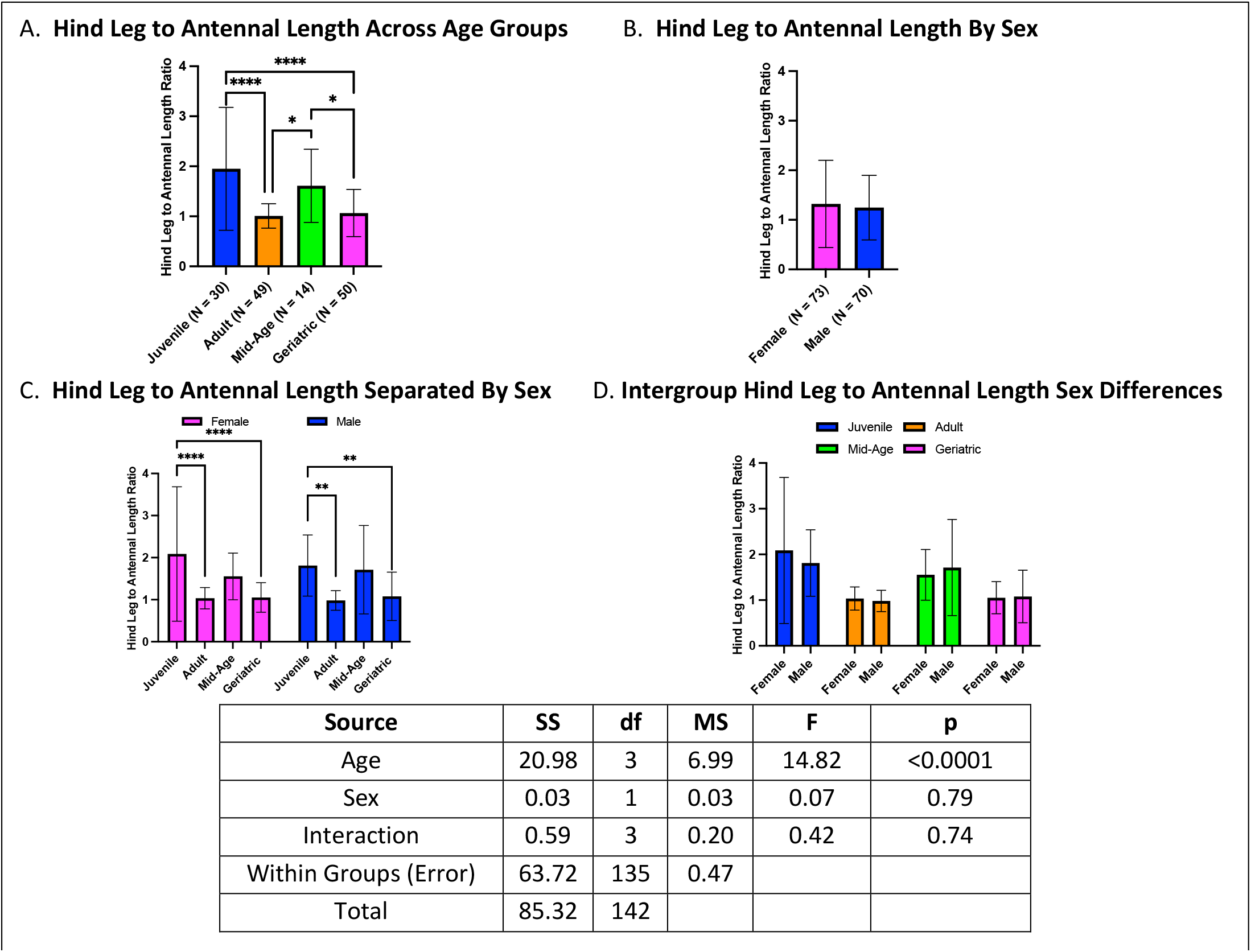
Hind leg to antennal length ratios across age and sex. **(A)** Adults and geriatric crickets had lower hindleg to antennal length ratios compared to juveniles and mid-age crickets. **(B)** No sex differences were observed in overall analysis. **(C)** When stratified by both age and sex, the same age-related trend persists in both males and females. **(D)** No sex differences were observed within any age group. (*p < 0.05, **p < 0.01, ***p < 0.001, ****p < 0.0001).

### Antennal length to weight ratios declines with age, with sex-specific differences in older cohorts

Antennal length to weight ratios exhibited declines with age, with geriatric crickets having lower ratios than adults, mid-age crickets having lower ratios than adults, and adults having higher ratios than juveniles (Figure 17A). A sex effect was observed, with males exhibiting higher antennal length to weight ratios than females in overall comparisons (Figure 17B). When stratified by age and sex, females exhibited a progressive decline with age, with mid-age and geriatric females having lower ratios than adults. In contrast, males exhibited a less gradual decline, with juveniles, mid-age, and geriatric crickets all having lower ratios than adults (Figure 17C). Within-group sex comparisons revealed that males had higher ratios than females in adult and geriatric cohorts, whereas no sex differences were observed in juveniles and mid-age crickets (Figure 17D).

**Figure 17.**
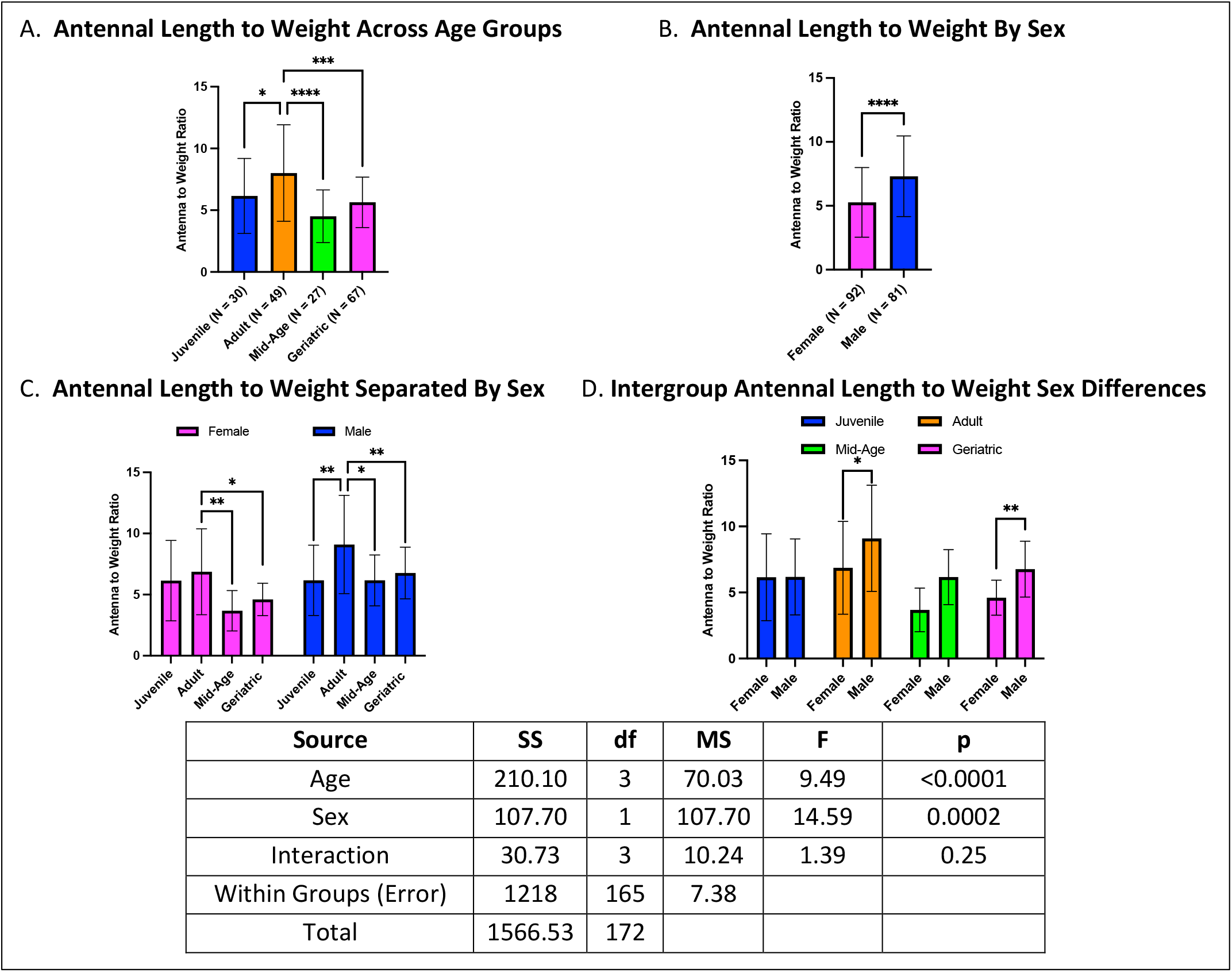
Antennal length to weight ratios across age and sex. **(A)** Geriatric and mid-age crickets exhibited lower antennal length to weight ratios than adults, with adults having higher ratios than juveniles. **(B)** Males had higher antennal length to weight ratios than females in overall comparisons. **(C)** Females exhibited a gradual decline in antennal length to weight ratios with age, whereas males showed a sharp decrease between adults and other age groups. **(D)** Sex differences were present in adults and geriatrics but not in juveniles or mid-age crickets. (*p < 0.05, **p < 0.01, ***p < 0.001, ****p < 0.0001).

## Discussion

Our findings provide novel insights into age- and sex-dependent morphological changes in house crickets, revealing distinct growth trajectories that influence locomotor function, sensory adaptation, and reproductive investment. Across multiple traits, we observed progressive increases in body weight, length, and appendage dimensions with age, with notable sex dimorphism emerging post-maturity. These results align with ecological and evolutionary theories of growth and resource allocation in insects [35–36].

Age-related increases in body weight, length, and appendage size indicate sustained somatic growth throughout adulthood, a pattern consistent with other hemimetabolous insects exhibiting prolonged post-maturity growth at diminishing rates [37]. Notably, geriatric crickets exhibited greater body dimensions than juveniles and adults but did not differ from mid-aged individuals, suggesting a plateau in late-life growth. The observed increases in femoral volume and CSA with age suggest structural reinforcement, potentially compensating for age-related declines in muscle efficiency [38–39]. As femoral volume serves as a proxy for structural mass, these changes may enhance muscle attachment sites for optimizing force regeneration during jumping [14,40] while simultaneously increasing load-bearing capacity to mitigate mechanical stress accumulation [41]. However, the observed decline in hind leg to body length and hind leg to weight ratios with age suggests disproportionate body mass increases relative to limb growth, potentially imposing biomechanical constraints on locomotion, as documented in other orthopterans [42]. Additionally, the inverse relationship between femoral SA/V ratio and age implies a reduction in relative surface area, which may impact heat dissipation and metabolic efficiency [43].

Sexual dimorphism was evident across multiple morphological traits, with females exhibiting greater body weight, length, and femoral dimensions than males, particularly in mid-aged and geriatric cohorts. This pattern aligns with established trends in orthopterans, where post-maturity sexual dimorphism emerges due to differential resource allocation [44–45]. Larger body size in females likely supports increased reproductive output, as enhanced body mass has been linked to greater fecundity and egg production [46–47]. In contrast, males maintained higher hind leg to body length and hind leg to weight ratios across adult and older age groups, suggesting a prioritization of locomotor efficiency, possibly as an adaptation for mate-searching and competitive behaviors [48]. The greater femoral volume observed in females may reflect biomechanical adaptations to support reproductive burdens [49], reinforcing the trade-offs between locomotion and reproductive investment.

Antenna to body length ratios increased with age, particularly in geriatric crickets, suggesting prolonged sensory investment that may compensate for declining locomotor capabilities [50]. While antennal length did not exhibit sex differences overall, sex-stratified analyses revealed distinct age-related trajectories, with males displaying a different growth pattern than females. The reduction in antennal length to weight ratio in aging crickets, particularly in females, suggests a shift in energetic allocation favoring reproductive or structural investments over sensory appendage growth.

Taken together, our findings highlight complex morphological adaptations influenced by aging and sex, with potential implications for locomotor performance, sensory function, and reproductive investment. The observed reductions in hind leg to body length ratios and femoral SA/V ratios suggest biomechanical trade-offs that may contribute to age-related declines in mobility. Future investigations should incorporate functional assays, including jump performance [51] and antennal-mediated navigation tasks such as the escape paradigm [52], to elucidate the physiological consequences of these morphological changes. Additionally, exploring the endocrine mechanisms underlying sexually dimorphic growth patterns may provide deeper insights into regulatory pathways governing aging in orthopterans.

To ensure the reproducibility and reliability of the house cricket as a model organism in aging and behavioral research, we implemented standardized husbandry practices, optimizing environmental conditions, housing densities, and dietary formulations. By minimizing confounding variables that may obscure biological signals, these refinements enhance data consistency and comparability across studies. Our refined husbandry protocol, incorporating self-determined photoperiods, co-housing both sexes, and carefully controlled diet and hydration strategies, establishes a framework for robust and reproducible research outcomes. Beyond fundamental husbandry, standardization in experimental design is critical to ensuring that observed physiological and behavioral changes are driven by biological processes rather than environmental variability. Mimicking natural habitats through varied lighting conditions, substrate availability, and microbiome exposure has been shown to enhance physiological resilience and behavioral fidelity, reinforcing the translational relevance of crickets as a model for studying aging and longevity. These refinements not only improve ecological validity but also provide a scalable system for future studies investigating age-related functional decline across species, such as cognitive decline (Scotto-Lomassese et al., 2003).

Despite the robustness of our findings, several limitations warrant consideration. First, while our study provides a comprehensive morphological assessment, we did not directly measure functional consequences such as locomotor performance, metabolic rates, or survival outcomes. Future research integrating behavioral assays and biomechanical testing will be critical for understanding how morphological changes influence organismal fitness. Second, environmental factors, including diet, temperature, and humidity, may impact growth trajectories. Although we controlled for these variables experimentally, natural populations may exhibit greater phenotypic variability. Another limitation pertains to the potential influence of behavioral plasticity on measured traits. Crickets exhibit phenotypic plasticity in response to environmental stimuli, and it remains unclear whether observed morphological differences reflect fixed developmental programs or are modulated by external factors such as competition or social interactions [53,54]. Longitudinal studies tracking individual crickets throughout their lifespan could disentangle intrinsic aging effects from extrinsic influences. Finally, our study did not assess femoral material properties, such as cuticular composition or density, which could provide further insight into age-related structural changes beyond morphological dimensions. Additionally, while we identified sex differences in femoral morphology, behavioral analyses are necessary to determine whether these differences translate to functional locomotor outcomes.

To address these limitations, we are integrating behavioral analyses into our study design, including assessments of locomotor performance, exploratory behavior, and cognitive function across age groups. Longitudinal tracking will allow for a finer-scale analysis of individual growth trajectories over time. Furthermore, we are expanding our investigation to include molecular markers of aging, which may provide deeper insight into the physiological mechanisms underlying observed morphological changes. Transcriptomic and proteomic approaches could further elucidate the genetic and biochemical pathways driving these changes, enhancing our understanding of aging and sexual dimorphism in house crickets.

Future refinements in cricket husbandry should explore the influence of microbiome diversity on lifespan and resilience to stress. Additionally, refining methodologies for raising crickets from eggs will be critical to maintaining controlled developmental conditions and minimizing cohort variability. The impact of oviposition substrate omission on reproductive health and feeding behavior warrants further investigation into its potential effects on metabolic regulation and stress responses. Advances in automated tracking technologies could further improve individual identification and behavioral monitoring, reducing handling stress while increasing data collection accuracy.

In conclusion, our study provides a comprehensive analysis of age- and sex-dependent morphological variation in house crickets, elucidating key developmental trends that influence body size, appendage length, and potential functional outcomes, as well as providing standardized husbandry practices essential for ensuring the reproducibility and reliability of the house cricket as a model organism in aging and behavioral research. With increasing interest in cricket models as translational systems for aging research [55], our findings contribute to a broader understanding of how aging and sexual dimorphism shape organismal structure. These insights have implications for studies of arthropod biomechanics, neurophysiology, and evolutionary ecology. By addressing current limitations and expanding our research focus, future work will further clarify the interplay between morphology, function, and environmental adaptation in aging crickets.

## Supporting information

Publishing licenses for Biorender figures

## References

[1] Kirkwood TB. Understanding the odd science of aging. Cell. 2005;120(4):437–447. doi:10.1016/j.cell.2005.01.027

[2] López-Otín C, Blasco MA, Partridge L, Serrano M, Kroemer G. The hallmarks of aging. Cell. 2013;153(6):1194–1217. doi:10.1016/j.cell.2013.05.039

[3] Liao GY, Pettan-Brewer C, Ladiges W. Comparison of age-related decline in C57BL/6J and CB6F1J male mice. PLoS One. 2024;19(12):e0306201. Published 2024 Dec 31. doi:10.1371/journal.pone.0306201

[4] Hendrickx JO, De Moudt S, Calus E, De Deyn PP, Van Dam D, De Meyer GRY. Age-related cognitive decline in spatial learning and memory of C57BL/6J mice. Behav Brain Res. 2022;418:113649. doi:10.1016/j.bbr.2021.113649

[5] Benice TS, Rizk A, Kohama S, Pfankuch T, Raber J. Sex-differences in age-related cognitive decline in C57BL/6J mice associated with increased brain microtubule-associated protein 2 and synaptophysin immunoreactivity. Neuroscience. 2006;137(2):413–423. doi:10.1016/j.neuroscience.2005.08.029

[6] Partridge L, Gems D. Mechanisms of ageing: public or private?. Nat Rev Genet. 2002;3(3):165–175. doi:10.1038/nrg753

[7] Tower J. Mitochondrial maintenance failure in aging and role of sexual dimorphism. Arch Biochem Biophys. 2015;576:17–31. doi:10.1016/j.abb.2014.10.008

[8] Finch CE, Tanzi RE. Genetics of aging. Science. 1997;278(5337):407–411. doi:10.1126/science.278.5337.407

[9] Chown SL, Gaston KJ. Body size variation in insects: a macroecological perspective. Biol Rev Camb Philos Soc. 2010;85(1):139–169. doi:10.1111/j.1469-185X.2009.00097.x

[10] Flatt T, Schmidt PS. Integrating evolutionary and molecular genetics of aging. Biochim Biophys Acta. 2009;1790(10):951–962. doi:10.1016/j.bbagen.2009.07.01

[11] Lyn JC, Naikkhwah W, Aksenov V, & Rollo CD. Influence of two methods of dietary restriction on life history features and aging of the cricket Acheta domesticus. Age (Dordr), 2011, 33(4):509–522.

[12] Liao GY, Rosenfeld M, Wezeman J, Ladiges W. The house cricket is an unrecognized but potentially powerful model for aging intervention studies. Aging Pathobiol Ther. 2024;6(1):39–41. doi:10.31491/apt.2024.03.135

[13] Scotto-Lomassese S, Strambi C, Strambi A, et al. Suppression of adult neurogenesis impairs olfactory learning and memory in an adult insect. J Neurosci. 2003;23(28):9289–9296. doi:10.1523/JNEUROSCI.23-28-09289.2003

[14] Burrows M, Morris O. Jumping and kicking in bush crickets. J Exp Biol. 2003;206(Pt 6):1035–1049. doi:10.1242/jeb.00214

[15] Mitchell SJ, MacArthur MR, Kane AE. Optimizing preclinical models of ageing for translation to clinical trials. Br J Clin Pharmacol. 2025;91(1):5–7. doi:10.1111/bcp.15902

[16] Mitchaothai J, Grabowski NT, Lertpatarakomol R, Trairatapiwan T, Lukkananukool A. Bacterial Contamination and Antimicrobial Resistance in Two-Spotted (Gryllus bimaculatus) and House (Acheta domesticus) Cricket Rearing and Harvesting Processes. Vet Sci. 2024;11(7):295. Published 2024 Jul 1. doi:10.3390/vetsci11070295

[17] Ponton F, Otálora-Luna F, Lefèvre T, et al. Water-seeking behavior in worm-infected crickets and reversibility of parasitic manipulation. Behav Ecol. 2011;22(2):392–400. doi:10.1093/beheco/arq215

[18] American Committee Of Medical Entomology American Society Of Tropical Medicine And Hygiene. Arthropod Containment Guidelines, Version 3.2. Vector Borne Zoonotic Dis. 2019;19(3):152–173. doi:10.1089/vbz.2018.2431

[19] Swallow J, Anderson D, Buckwell AC, et al. Guidance on the transport of laboratory animals. Lab Anim. 2005;39(1):1–39. doi:10.1258/0023677052886493

[20] Sensini, F., Inta, D., Palme, R., Brandwein, C., Pfeiffer, N., Riva, M. A., Gass, P., & Mallien, A. S. (2020). The impact of handling technique and handling frequency on laboratory mouse welfare is sex-specific. Scientific reports, 10(1), 17281. 10.1038/s41598-020-74279-3

[21] Gray DA. Female house crickets, Acheta domesticus, prefer the chirps of large males. Anim Behav. 1997;54(6):1553–1562. doi:10.1006/anbe.1997.0584

[22] Miller, J. P., Krueger, S., Heys, J. J., & Gedeon, T. (2011). Quantitative characterization of the filiform mechanosensory hair array on the cricket cercus. PloS one, 6(11), e27873. 10.1371/journal.pone.0027873

[23] Hochkirch, A., & Gröning, J. (2008). Sexual size dimorphism in Orthoptera (sensu stricto)—A review. Journal of Orthoptera Research, 17(2), 189–196. 10.1665/1082-6467-17.2.189

[24] Fluker’s Cricket Biology Guide. Fluker’s Cricket Farms. Accessed February 26, 2025. https://flukerfarms.com/content/Cricket.pdf.

[25] Ventura MK, Stull VJ, Paskewitz SM. Maintaining Laboratory Cultures of Gryllus bimaculatus, a Versatile Orthopteran Model for Insect Agriculture and Invertebrate Physiology. J Vis Exp. 2022;(184):10.3791/63277. Published 2022 Jun 8. doi:10.3791/63277

[26] Clifford CW, Roe RM, & Woodring J. Rearing methods for obtaining house crickets, Acheta domesticus, of known age, sex, and instar. Annals of the Entomological Society of America, 1977, 70(1):69–74.

[27] Levy K, Barnea A, Tauber E, Ayali A. Crickets in the spotlight: exploring the impact of light on circadian behavior. J Comp Physiol A Neuroethol Sens Neural Behav Physiol. 2024;210(2):267–279. doi:10.1007/s00359-023-01686-y

[28] Ghosal K, Gupta M, Killian KA. Agonistic behavior enhances adult neurogenesis in male Acheta domesticus crickets. J Exp Biol. 2009;212(Pt 13):2045-2056. doi:10.1242/jeb.026682

[29] Takacs J, Bryon A, Jensen AB, van Loon JJA, Ros VID. Effects of Temperature and Density on House Cricket Survival and Growth and on the Prevalence of Acheta Domesticus Densovirus. Insects. 2023;14(7):588. Published 2023 Jun 29. doi:10.3390/insects14070588

[30] Gutiérrez Y, Fresch M, Ott D, Brockmeyer J, Scherber C. Diet composition and social environment determine food consumption, phenotype and fecundity in an omnivorous insect. R Soc Open Sci. 2020;7(4):200100. Published 2020 Apr 22. doi:10.1098/rsos.200100

[31] Murtaugh MP & Denlinger DL. Physiological regulation of long-term oviposition in the house cricket, Acheta domesticus. J Insect Physiol. 1985;31(8):611–617. doi:10.1016/0022-1910(85)90059-9

[32] Dobson GP, Letson HL, Biros E, Morris J. Specific pathogen-free (SPF) animal status as a variable in biomedical research: Have we come full circle?. EBioMedicine. 2019;41:42–43. doi:10.1016/j.ebiom.2019.02.038

[33] Masopust D, Sivula CP, Jameson SC. Of Mice, Dirty Mice, and Men: Using Mice To Understand Human Immunology. J Immunol. 2017;199(2):383–388. doi:10.4049/jimmunol.1700453

[34] Leary S. Guidelines for the euthanasia of animals. American Veterinary Medical Association. 2020. Accessed February 25, 2025. https://www.avma.org/resources-tools/avma-policies/avma-guidelines-euthanasia-animals.

[35] Nijhout HF, Emlen DJ. Competition among body parts in the development and evolution of insect morphology. Proc Natl Acad Sci U S A. 1998;95(7):3685–3689. doi:10.1073/pnas.95.7.3685

[36] Stillwell RC, Blanckenhorn WU, Teder T, Davidowitz G, Fox CW. Sex differences in phenotypic plasticity affect variation in sexual size dimorphism in insects: from physiology to evolution. Annu Rev Entomol. 2010;55:227–245. doi:10.1146/annurev-ento-112408-085500

[37] Chapman RF. The Insects Structure and Function Ed by R.F. Chapman. Cambridge Univ. Press; 2013.

[38] Katz, SL, Gosline, JM. Ontogenetic Scaling of Jump Performance in the African Desert Locust (Schistocerca Gregaria). J Exp Biol. 1994;177(1):81–111. doi:10.1242/jeb.177.1.81

[39] Huie JM, Summers AP, Kawano SM. SegmentGeometry: A Tool for Measuring Second Moment of Area in 3D Slicer. Integr Org Biol. 2022;4(1):obac009. Published 2022 Feb 28. doi:10.1093/iob/obac009

[40] Burrows M. Do the enlarged hind legs of male thick-legged flower beetles contribute to take-off or mating?. J Exp Biol. 2020;223(Pt 1):jeb212670. Published 2020 Jan 2. doi:10.1242/jeb.212670

[41] Douglas W. Whitman “The significance of body size in the Orthoptera: a review,” Journal of Orthoptera Research, 17(2), 117–134, (1 December 2008)

[42] Gabriel, JM. The Development of the Locust Jumping Mechanism: 1. Allometric Growth and its Effect on Jumping Performance. J Exp Biol. 1985;118(1):313–326. doi:10.1242/jeb.118.1.313

[43] Schmidt-Nielsen, K. Scaling, why is animal size so important. Cambridge Univ. Press; 1984.

[44] Honěk, A. Intraspecific Variation in Body Size and Fecundity in Insects: A General Relationship. Oikos 1993;66(3):483–492. 10.2307/3544943.

[45] Bonduriansky R. Sexual selection and allometry: a critical reappraisal of the evidence and ideas. Evolution. 2007;61(4):838–849. doi:10.1111/j.1558-5646.2007.00081.x

[46] Savage VM, Gilloly JF, Brown JH, Charnov EL. Effects of body size and temperature on population growth. Am Nat. 2004;163(3):429–441. doi:10.1086/381872

[47] Bonduriansky R. The evolution of male mate choice in insects: a synthesis of ideas and evidence. Biol Rev Camb Philos Soc. 2001;76(3):305–339. doi:10.1017/s1464793101005693

[48] Boisseau RP, Büscher TH, Klawitter LJ, Gorb SN, Emlen DJ, Tobalske BW. Multi-modal locomotor costs favor smaller males in a sexually dimorphic leaf-mimicking insect. BMC Ecol Evol. 2022;22(1):39. Published 2022 Mar 28. doi:10.1186/s12862-022-01993-z

[49] Bertram, S.M., Thomson, I.R., Auguste, B.L., Dawson, J.W., & Darveau, C. Variation in cricket acoustic mate attraction signalling explained by body morphology and metabolic differences. Animal Behaviour. 2011;82; 1255–1261. doi:10.1016/j.anbehav.2011.08.021

[50] Gronenberg W, Heeren S, HÖLldobler B. Age-dependent and task-related morphological changes in the brain and the mushroom bodies of the ant Camponotus floridanus. J Exp Biol. 1996;199(Pt 9):2011–2019. doi:10.1242/jeb.199.9.2011

[51] Lachenicht MW, Clusella-Trullas S, Boardman L, Le Roux C, Terblanche JS. Effects of acclimation temperature on thermal tolerance, locomotion performance and respiratory metabolism in Acheta domesticus L. (Orthoptera: Gryllidae). J Insect Physiol. 2010;56(7):822–830. doi:10.1016/j.jinsphys.2010.02.010

[52] Scotto-Lomassese S, Strambi C, Strambi A, et al. Suppression of adult neurogenesis impairs olfactory learning and memory in an adult insect. J Neurosci. 2003;23(28):9289–9296. doi:10.1523/JNEUROSCI.23-28-09289.2003

[53] Moczek AP. Phenotypic plasticity and diversity in insects. Philos Trans R Soc Lond B Biol Sci. 2010;365(1540):593–603. doi:10.1098/rstb.2009.0263

[54] Corona M, Libbrecht R, Wheeler DE. Molecular mechanisms of phenotypic plasticity in social insects. Curr Opin Insect Sci. 2016;13:55–60. doi:10.1016/j.cois.2015.12.003

[55] Mito T, Ishimaru Y, Watanabe T, et al. Cricket: The third domesticated insect. Curr Top Dev Biol. 2022;147:291–306. doi:10.1016/bs.ctdb.2022.02.003

