## Supplementary material for "Morphological features of the domestic house cricket (*Acheta domesticus*) for translational aging studies": Publishing licenses for Biorender figures: 4 - Publication License Mar-26-2025.pdf

BIO RENDER

49 Spadina Ave. Suite 200
Toronto ON M5V 2J1 Canada
www.biorender.com

### Confirmation of Publication and Licensing Rights - Open Access

**March 26th, 2025**

***Subscription Type:*** *Student Plan - Academic*
***Agreement number:*** *DJ282O8A2U*
***Publisher Name:*** *bioRxiv*

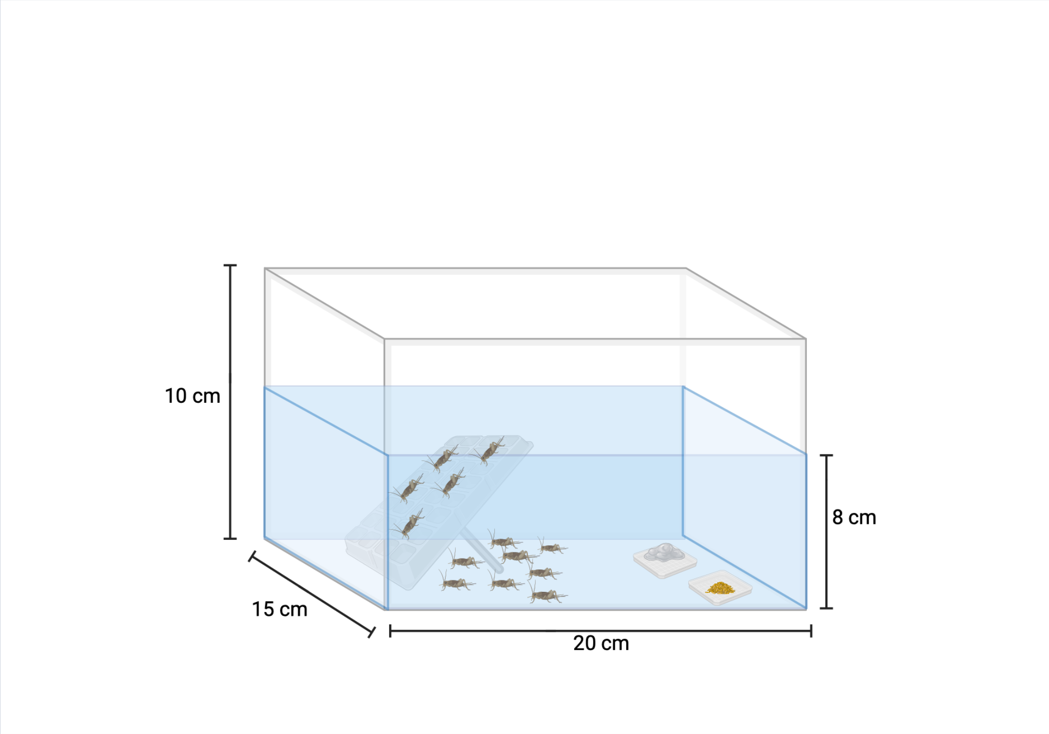


*For any questions regarding this document, or other questions about publishing with BioRender, please refer to our BioRender Publication Guide, or contact BioRender Support at.*
