## Supplementary material for "Morphological features of the domestic house cricket (*Acheta domesticus*) for translational aging studies": Publishing licenses for Biorender figures: 5 - Publication License Mar-26-2025.pdf

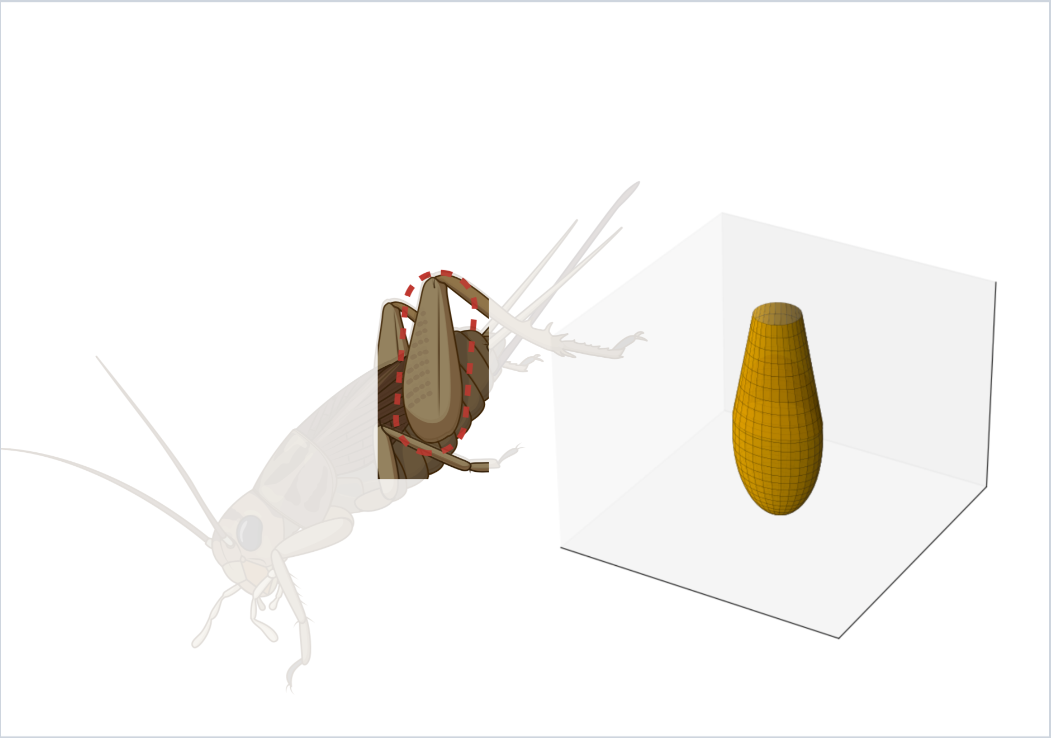

*For any questions regarding this document, or other questions about publishing with BioRender, please refer to our BioRender Publication Guide, or contact BioRender Support at.*
